# Patient-specific responses to *SMN2* splice-modifying treatments in spinal muscular atrophy fibroblasts

**DOI:** 10.1101/2024.07.24.604892

**Authors:** Ilaria Signoria, Maria M. Zwartkruis, Lotte Geerlofs, Elena Perenthaler, Kiterie M.E. Faller, Rachel James, Harriet McHale-Owen, Jared W. Green, Joris Kortooms, Sophie H. Snellen, Fay-Lynn Asselman, Thomas H. Gillingwater, Gabriella Viero, Renske I. Wadman, W. Ludo van der Pol, Ewout J.N. Groen

## Abstract

The availability of three therapies for the neuromuscular disease spinal muscular atrophy (SMA) highlights the need to match patients to the optimal treatment. Two of these treatments (nusinersen and risdiplam) target splicing of *SMN2*, but treatment outcomes vary from patient to patient. An incomplete understanding of the complex interactions between SMA genetics, SMN protein and mRNA levels, and gene-targeting treatments, limits our ability to explain this variability and identify optimal treatment strategies for individual patients. To address this, we analyzed responses to nusinersen and risdiplam in 45 primary fibroblast cell lines. Pre-treatment *SMN2-FL*, *SMN2Δ7* mRNA, and SMN protein levels were influenced by *SMN2* copy number, age, and sex. After treatment, SMN and mRNA levels were more heterogeneous. In 43% of patients, response to both therapies was similar, but in 57% one treatment led to a significantly higher SMN increase than the other treatment. Younger age, higher *SMN2* copy number, and higher SMN levels before treatment predicted better *in vitro* efficacy. These findings showcase patient-derived fibroblasts as a tool for identifying molecular predictors for personalized treatment.

## Introduction

The increasing availability of gene and gene-targeting therapies offers significant hope for patients with genetic diseases. In neurology, spinal muscular atrophy (SMA) leads these advancements, with three gene-targeting treatments approved for use and over a decade of follow-up since the earliest clinical trials (Chaytow *et al*, 2021). SMA, a severe childhood-onset neuromuscular disease, is characterized by the degeneration of spinal motor neurons, resulting in progressive weakness, respiratory insufficiency, and often premature death (Mercuri *et al*, 2022). SMA is caused by homozygous loss-of-function of *SMN1*, leading to insufficient SMN protein levels. The severity of SMA is influenced by the number of copies of the partially functional *SMN2* gene (Lefebvre *et al*, 1995; Wadman *et al*, 2017; Calucho *et al*, 2018). A C-to-T transition in exon 7 of *SMN2* mostly results in the production of truncated *SMN2Δ7* mRNA, which is translated to an unstable and dysfunctional SMNΔ7 protein (Monani *et al*, 1999; Lorson *et al*, 1999, 1). Low levels of full-length *SMN2* (*SMN2-FL*) mRNA and SMN protein are also produced from each *SMN2* copy, creating a negative correlation between *SMN2* copy number, SMN protein levels, and SMA severity (Monani *et al*, 1999; Lorson *et al*, 1999). SMN mRNA and protein levels, and *SMN2* splicing, vary depending on age and developmental stage, and cell- or tissue type (Kobayashi *et al*, 2011; Groen *et al*, 2018; Wadman *et al*, 2016; Ramos *et al*, 2019; Zaworski *et al*, 2016). The relationship between *SMN2* copy number, *SMN2-FL* and *SMN2Δ7* mRNA, and SMN protein levels, however, is complex and remains incompletely understood.

Currently available treatments for SMA are the gene replacement therapy onasemnogene abeparvovec-xioi (Zolgensma), an adeno-associated virus serotype 9 (AAV9) containing the *SMN* open reading frame (ORF) (Mendell *et al*, 2017; Valori *et al*, 2010); and two *SMN2* splice-modifiers, nusinersen (Spinraza, an antisense oligonucleotide delivered via intrathecal injections) (Finkel *et al*, 2017; Mercuri *et al*, 2018; Hua *et al*, 2008) and risdiplam (Evrysdi, a daily oral small molecule) (Naryshkin *et al*, 2014; Baranello *et al*, 2021). Since SMA treatments are most effective when started presymptomatically (Strauss *et al*, 2022a, 2022b), newborns are now commonly screened for SMA. Early genetic diagnosis and treatment often lead to spectacular improvements in survival, motor function, and quality of life compared to the natural history of SMA (Strauss *et al*, 2022a, 2022b; Crawford *et al*, 2023). Around the world, however, 10,000s of SMA patients were already symptomatic as gene-targeting treatments for SMA became available. Because the use of Zolgensma is limited to 2 years of age, symptomatic children and adults often received one of the *SMN2* splice-modifiers. They still benefit from these treatments, but treatment effects are more modest and variable. A meta-analysis of real-world studies on nusinersen illustrated that outcomes vary significantly: 10% of patients experiencing a further decline, while others achieved new motor milestones or stabilized (Coratti *et al*, 2021). For risdiplam, fewer real-world studies exist, but clinical trials suggest similar effectiveness and variability (Baranello *et al*, 2021; Oskoui *et al*, 2023). No head-to-head trials between nusinersen and risdiplam have been conducted, complicating optimal treatment selection for individual patients (Kokaliaris *et al*, 2024). Variation in patient age, *SMN2* copy number, disease duration at treatment start, and comorbidities complicate comparisons and understanding of treatment outcomes, highlighting ongoing challenges for patients and their families, especially when treatment started symptomatically.

The direct interaction of *SMN2* splice-modifying treatments with *SMN2* pre-mRNA (Singh *et al*, 2017, 2020) highlights the need to better understand the relationship between *SMN2* copy number, *SMN2*-derived mRNA levels and SMN protein expression. This improved understanding would help explain the variability in treatment outcomes and support more informed treatment decisions. Primary fibroblasts are particularly suited to address this issue: they preserve genetic and epigenetic signatures of the donor (Sturm *et al*, 2019), whilst being homogeneous and scalable (Kumbier *et al*, 2024), thus enabling personalized studies across many patients. We here therefore used 45 primary fibroblast cell lines (10 from healthy donors, 35 from patients with SMA) to characterize *in vitro* response to *SMN2* splice-modifying treatments. Using this approach, we found that SMN levels before treatment were influenced by *SMN2* copy number, age and sex. After treatment, SMN mRNA and protein levels were significantly more heterogeneous and younger age, higher *SMN2* copy number and higher pre-treatment SMN protein levels were the main predictors of *in vitro* treatment efficacy. Importantly, 57% of the cell lines investigated showed a preferred response to one of the treatments, despite similar treatment mechanisms. Our findings emphasize the effectiveness of patient-derived primary fibroblasts in modelling and predicting outcomes after exposure to SMN-restoring therapies and identifying predictors of treatment efficacy in SMA.

## Results

### Patient-derived primary fibroblasts reflect key molecular characteristics of SMA

We first characterized a large cohort of representative, untreated primary fibroblasts from our biobank that were selected based on age, SMA type, sex and *SMN2* copy number. We included cell lines from 35 SMA patients and 10 healthy donors, hereafter referred to as control cell lines (**Material and methods** and **Supplementary table 2**). As working with many primary cell lines requires collection and analysis over a prolonged period, we developed a standardized tissue culture and analysis pipeline (**Fig. S1**). Within this standardized pipeline, we established the effect of passage number, confluency, cell growth, lipofection, cell batch and culture time on SMN levels (**Fig. S2**), allowing us to reliably and reproducibly determine SMN expression over time and control for variability between different experiments.

We measured *SMN1*, *SMN2-FL* and *SMN2Δ7* mRNA levels along with SMN protein expression in each of the fibroblast cell lines using droplet digital PCR (ddPCR) and semi-quantitative western blotting (**Fig. S2I** and as described previously (Groen *et al*, 2018; Wadman *et al*, 2020)), and cellular morphology using fluoresecent microscopy (**Fig. S3A**). As expected, only control fibroblasts expressed *SMN1* mRNA (**Fig. 1A**). *SMN2-FL* mRNA expression was dependent on *SMN2* copy number (*P* = 6.84e-08), where cell lines with fewer *SMN2* copies expressed less *SMN2-FL* than fibroblasts with a higher number of *SMN2* copies (**Fig. 1B**). Similarly, we observed that *SMN2Δ7* mRNA levels (**Fig. 1C**) also depend on *SMN2* copy number (*P* = 7.16e-10) and cell lines with fewer *SMN2* copies expressed less *SMN2Δ7*. We found that *SMN2-FL* made up 51% of total *SMN2* mRNA, a distribution that was comparable between control and SMA fibroblasts (**Fig. S4A** and **S4B**). Moreover, we observed a significant correlation between *SMN2-FL* and *SMN2Δ7* levels (**Fig. S4C**, R^2^ = 0.41, *P* = 3e-05).

**Figure 1.**
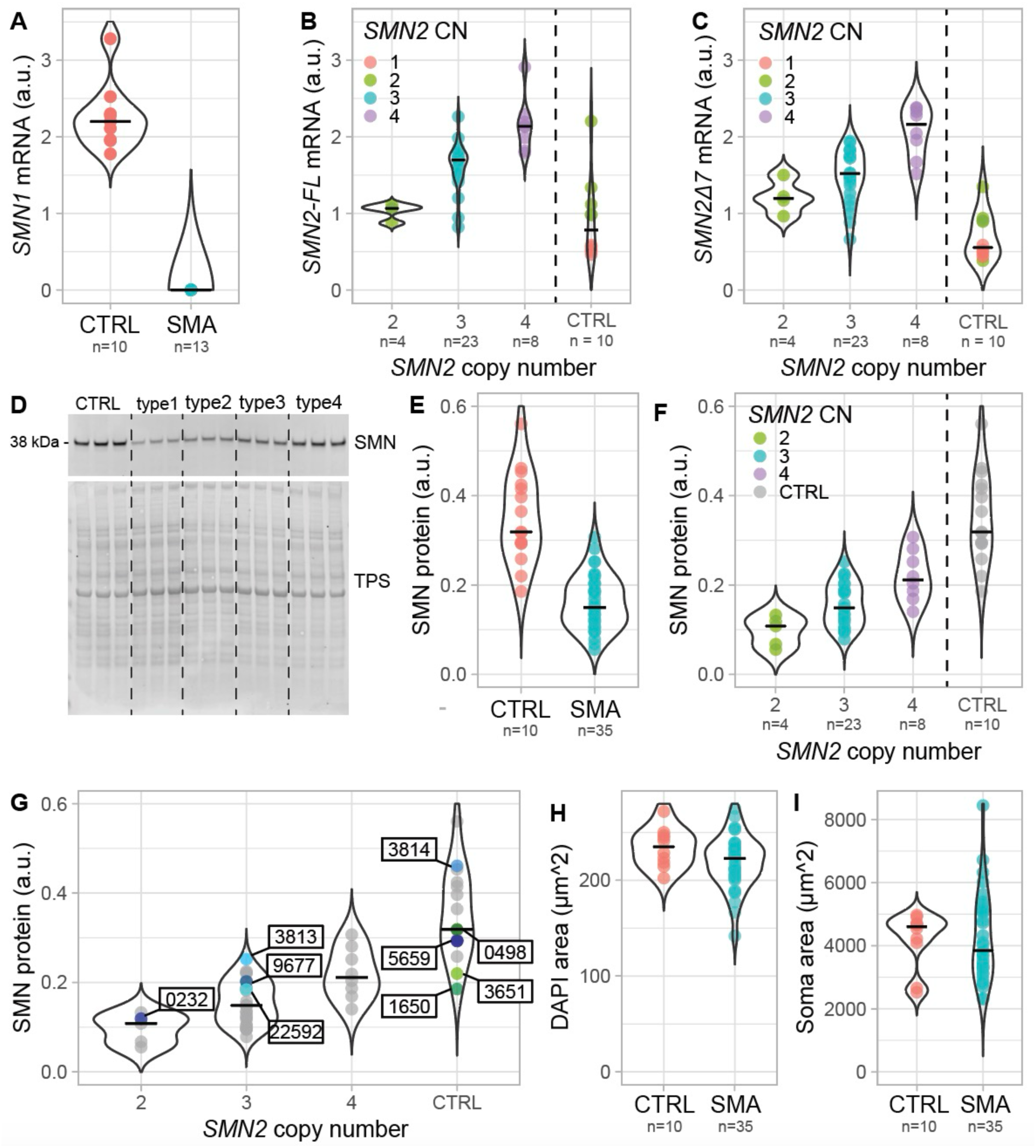
Patient-derived primary fibroblasts reflect key molecular characteristics and model heterogeneity observed in SMA. **(A)** *SMN1* mRNA expression in control (n=10) and SMA (n=13) patient-derived primary fibroblasts. Each dot corresponds to the average of technical triplicate of each cell line. **(B)** *SMN2-FL* expression in control (n=10) and SMA patient-derived primary fibroblasts with 2x *SMN2* copies (n=4), 3x *SMN2* copies (n=23) and 4x *SMN2* copies (n=8). *SMN2-FL* mRNA expression in SMA patient derived primary fibroblasts depends on *SMN2* copy number (one-way ANOVA *P* = 6.84e-08, 2 vs. 3 *P* = 0.03, 2 vs. 4 *P* = 0.001, 3 vs. 4 *P* = 0.005). Each dot corresponds to the average of technical triplicates for each cell line. **(C)** *SMN2Δ7* mRNA expression in control (n=10) and SMA patient-derived primary fibroblasts with 2x *SMN2* copies (n=4), 3x *SMN2* copies (n=23) and 4x *SMN2* copies (n=8). *SMN2Δ7* mRNA expression depends on *SMN2* copy number (one-way ANOVA *P* = 7.16e-10, 2 vs. 4 *P* = 0.001, 3 vs. 4 *P* = 0.001). Each dot corresponds to the average of technical triplicate of each cell line. **(D)** Representative western blot of SMN protein in control and SMA type 1, type 2, type 3 and type 4 fibroblasts. **(E)** Normalized SMN protein expression levels in control (n=10) and SMA (n=35) patient-derived primary fibroblasts. Control cell lines express statistically significant higher amount of SMN protein (Welch two sample t-test, *P* = 3e-05). Each dot corresponds to the average of a technical triplicate for each cell line. **(F)** Normalized SMN protein expression levels in control (n=10) and SMA fibroblasts with 2x *SMN2* copies (n=4), 3x *SMN2* copies (n=23) and 4x *SMN2* copies (n=8). SMN protein expression shows a partial dependency on *SMN2* copy number (one-way ANOVA, *P* = 2e-04, 2 vs. 4 *P* = 3e-04, 3 vs. 4 *P* = 0.003). Each dot corresponds to the average of a technical triplicate for each cell line. **(G)** Normalized SMN protein expression levels in Coriell (GM00232, GM03813, GM09677, GM22591, GM03814, GM00498, GM05659, GM03651, GM01650) cell lines compared to cell lines from our biobank (as in **(F)**, depicted in grey). Each dot corresponds to the average of technical triplicate of each cell line. **(H)** DAPI area (µm²) of control (n=10) and SMA (n=35) patient-derived primary fibroblasts (Welch two Sample t-test, *P* = 0.07). Each dot corresponds to the average nuclear size of each cell line. **(I)** F-actin (soma) area of control (n=10) and SMA (n=35) patient-derived primary fibroblasts (Welch two Sample t-test, *P* = 0.88). Each dot corresponds to the average soma size of each cell line. a.u. = arbitrary unit; CN = copy number.

At the protein level, SMA-derived fibroblasts expressed 2.4-fold lower levels of SMN protein compared to control fibroblasts (*P* = 1.7e-11, **Fig. 1D** and **1E**). SMN protein expression was dependent on *SMN2* copy number (**Fig. 1F**, *P* = 2e-04) and cells with lower copy number expressed less SMN. The correlation between *SMN2-FL* and *SMN2Δ7* mRNA levels, and protein expression was significant but limited (**Fig. S5A** and **S5B**; R^2^ = 0.2, *P* = 0.007 and R^2^ = 0.015, *P* = 0.02, respectively), suggesting the presence of other, unknown factors that regulate *SMN2-FL-*translation or degradation. As previous studies using fibroblasts often included cells from the Coriell repository (e.g. (Brown *et al*, 2022; James *et al*, 2024; Kordala *et al*, 2023)), we determined SMN expression in four SMA and five control Coriell cell lines and compared them to cell lines from our biobank (**Fig. 1G**). Overall, SMN expression in the commonly-used Coriell cells was comparable to cells from our biobank, although Coriell cell lines with 3x *SMN2* had SMN levels that were higher than typical for the genotype (*P* = 0.03). Finally, we analysed cellular morphology by measuring the size of the nucleus and soma for each cell line (**Fig. 1H-I**). Although the area of both the nucleus and the cell soma varied considerably between and within cell lines (**Fig. S3B-E**), we did not identify any consistent, statistically significant differences for either morphological variable between SMA and control cell lines. We therefore focused on *SMN2-FL/Δ7* mRNA and SMN protein as readouts for further experiments.

### Younger age and male sex influence SMN mRNA and protein expression

To explore which patient characteristics influenced SMN mRNA and protein levels, we compared them to SMA type, sex and age. We found no clear relationship between SMA type and levels of *SMN2* mRNA or SMN protein (**Fig. S4F**, **S4G**, and **S5D**). The only significant differences were observed between type 1b and other types and were primarily driven by *SMN2* copy number. We did not observe a difference in SMN expression between male and female patients (*P* = 0.28). However, stratifying by sex revealed an enhanced correlation between *SMN2-FL* mRNA and SMN protein levels in male patients (R^2^ = 0.37, *P* = 0.005), while this correlation was absent in females (**Fig. S5E**, R^2^ = 0.00048, *P* = 0.94). Notably, males and females had unequal numbers of cells with 4x SMN2 copies, which may partially explain these results. Finally, we investigated whether *SMN2-FL* or *SMN2Δ7* mRNA and SMN protein expression in fibroblasts were influenced by age (**Fig. S4D, S4E** and **S5C**). For mRNA, we found a correlation between *SMN2Δ7* expression and age in children (<18 years, R^2^ = 0.44, *P* = 0.00082) but not in adults (R^2^ = 0.013, *P* = 0.71). For protein, we similarly observed a positive correlation between SMN expression and age in children (<18 years, R^2^ = 0.49, *P* = 0.0003) but not adults (R^2^ = 0.02, *P* = 0.65). Overall, the effect of patient characteristics other than *SMN2* copy number on SMN protein and mRNA levels was limited in untreated cells.

### SMN2 splice-modifiers increase in vitro variability of SMN expression

We next treated cells with the *SMN2* splice-modifying drugs nusinersen or risdiplam. We observed a complete depletion of *SMN2Δ7* mRNA after both treatments, a ∼2 to 2.5 fold increase in *SMN2-FL* mRNA in SMA (nusinersen *P* = 6.9e-15, risdiplam *P* = 2e-16) and a ∼2 fold increase in control fibroblasts (nusinersen *P* = 0.001, risdiplam *P* = 0.0002, **Fig. 2A** and **2B**). Nusinersen resulted in higher *SMN2-FL* mRNA levels after treatment than risdiplam in SMA fibroblasts (*P* = 0.0007, **Fig. 2B**), so *SMN2-FL* mRNA levels from the same cell lines after treatment correlated incompletely (R^2^ = 0.39, *P* = 6.5e-05, **Fig. S6A**). At the protein level, a significant increase in SMN was observed across all SMA cell lines (nusinersen: *P* = 1.1e-10, risdiplam: *P* = 4.1e-10, **Fig. 2C** and **2D**), with no difference between the two treatments (R^2^ = 0.83, *P* = 4.3e-14, **Fig. S6B**). In addition to increasing SMN levels, both treatments also increased the variability of both *SMN2-FL* mRNA (standard deviation of untreated vs. nusinersen *P* = 4.0e-09, and vs. risdiplam *P* = 6.2e-05) and SMN protein expression between cell lines (standard deviation of untreated vs. nusinersen *P* = 4.0e-04, and vs. risdiplam *P* = 2.0e-4). Although *SMN2Δ7* mRNA was always completely depleted after treatment, this did not lead to a consistent increase in levels of *SMN2-FL* mRNA after treatment, illustrated by a limited correlation between *SMN2Δ7* levels before and *SMN2-FL* levels after treatment (nusinersen: R^2^ = 0.43, *P* = 6.6e-07; risdiplam: R^2^ = 0.48, *P* = 1.0e-07, **Fig. S6C**). We found no correlation between SMN protein relative change and *SMN2-FL* mRNA relative change after treatment (**Fig. S6E**), suggesting that - similarly to what we previously observed in untreated cells (**Fig. S5A**) – other, unknown factors play an important role in regulating SMN protein levels.

**Figure 2.**
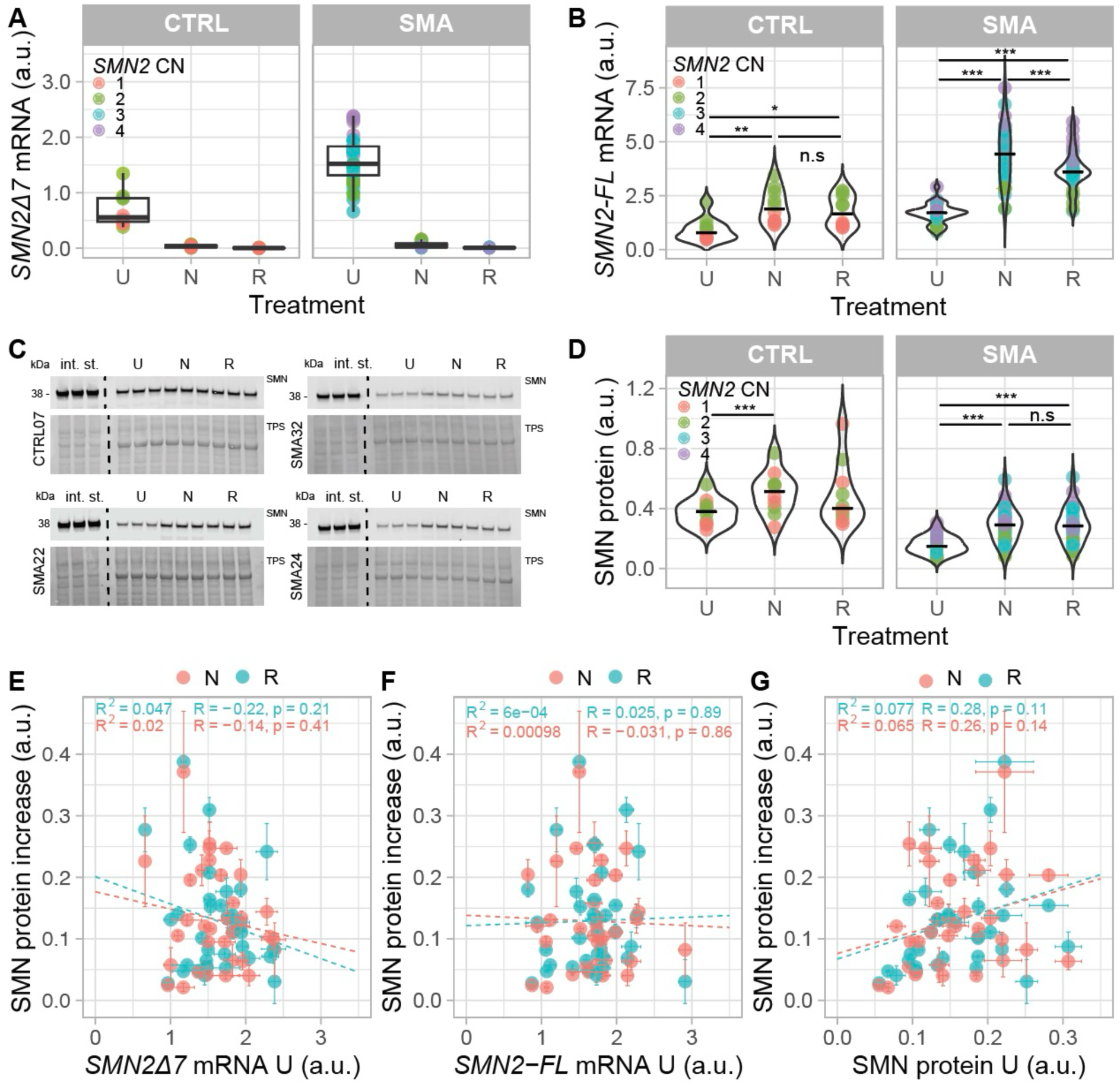
Effect of *SMN2* splice-modifying treatments on *SMN2-FL*, *SMN2Δ7* and SMN expression levels in control and SMA patient-derived primary fibroblasts. **(A)** *SMN2Δ7* mRNA expression levels in control (n=10) and SMA (n=35) patient-derived primary fibroblasts before and after *in vitro* treatment with either nusinersen (N) or risdiplam (R). *SMN2Δ7* mRNA was completely depleted after treatment in both control (paired Welch t-test, Bonferroni correction of p-values U vs. N *P* = 0.00029; U vs. R *P* = 0.00016) and SMA primary fibroblasts (paired Welch t-test, Bonferroni correction of p-values U vs. N *P* < 2e-16; U vs. R *P* < 2e-16). Each dot corresponds to the average of a technical triplicate of each cell line. **(B)** *SMN2-FL* mRNA expression levels in control (n=10) and SMA (n=35) patient-derived primary fibroblasts before and after *in vitro* treatment with either nusinersen (N) or risdiplam (R). *SMN2-FL* RNA increases after treatment in control (paired Welch t-test, Bonferroni correction of p-values U vs. N *P* = 0.001; U vs. R *P* = 0.0002) and SMA fibroblasts (paired Welch t-test, Bonferroni correction of p-values U vs. N *P* = 6.9e-15; U vs. R *P* = 2e-16). There is a significant difference in *SMN2-FL* levels between nusinersen and risdiplam treatment in SMA (paired Welch t-test, Bonferroni correction of p-values R vs. N *P* = 0.0007) but not in control fibroblasts (paired Welch t-test, Bonferroni correction of p-values U vs. N *P* = 0.12). Each dot corresponds to the average of a technical triplicate of each cell line. **(C)** Representative western blot of SMN protein expression in four different SMA fibroblast cell lines untreated (U) and after *in vitro* treatment with either nusinersen (N) or risdiplam (R). **(D)** Normalized expression levels in control (n=10) and SMA (n=35) patient-derived primary fibroblasts untreated (U) and after treatment with either nusinersen (N) or risdiplam (R). SMN protein significantly increases after treatment in SMA primary fibroblasts (paired Welch t-test, Bonferroni correction of p-values U vs. N *P* = 1.1e-10; U vs. R *P* = 4.1e-10) but not in control primary fibroblasts treated with risdiplam (paired Welch t-test, Bonferroni correction of p-values U vs. N *P* = 0.001; U vs. R *P* = 0.16). There is no significant difference in SMN protein increases between nusinersen and risdiplam treatment in both control and SMA patient-derived primary fibroblasts (paired Welch t-test, Bonferroni correction of p-values control N vs. R *P* = 1; SMA N vs. R *P* = 1). Each dot corresponds to the average of the technical triplicate of each cell line. **(E)** Relationship between *SMN2Δ7* mRNA in untreated cells (U) and SMN protein increase levels in SMA patient-derived primary fibroblasts after *in vitro* treatment with risdiplam (R, light blue) or nusinersen (N, coral). Data are represented as the average of the technical triplicate ± standard deviation. Regression line (dashed line), Pearson correlation coefficient (R), p-value (p) and the coefficient of determination (R²) are displayed. **(F)** Relationship between *SMN2-FL* mRNA in untreated cells (U) and SMN protein increase levels SMA patient-derived primary fibroblasts after *in vitro* treatment with risdiplam (R, light blue) or nusinersen (N, coral). Data are represented as the average of the technical triplicate ± standard deviation. Regression line (dashed line), Pearson correlation coefficient (R), p-value (p) and the coefficient of determination (R²) are displayed. **(G)** Relationship between SMN protein in untreated cells (U) and SMN protein increase in SMA patient-derived primary fibroblasts after *in vitro* treatment with risdiplam (R, light blue) or nusinersen (N, coral). Data are represented as the average of the technical triplicate ± standard deviation. Regression line (dashed line), Pearson correlation coefficient (R), p-value (p) and the coefficient of determination (R²) are displayed. TPS = total protein staining; in. st = internal standard; U = untreated; N = nusinersen; R = risdiplam; * = *P* < 0.05.; * = *P* < 0.01; * = *P* < 0.001; a.u. = arbitrary unit; CN = copy number.

Strikingly, as both nusinersen and risdiplam promote *SMN2* exon 7 inclusion, we hypothesized that the potential to increase SMN protein post-treatment would mostly be dependent on *SMN2Δ7* mRNA levels before treatment but found no correlation (**Fig. 2E**). Similarly, *SMN2-FL* mRNA and SMN protein expression before treatment did not correlate with the observed SMN increase after treatment (**Fig. 2F** and **2G**). However, when assessing total levels of SMN protein – rather than its increase after treatment – we found a correlation between SMN expression before and after treatment (nusinersen: R^2^ = 0.51, *P* = 1.3e-06; risdiplam: R^2^ = 0.51, *P* = 1.3e-06, **Fig. S6D**), suggesting the presence of patient-specific factors that regulate overall SMN protein expression that are not directly influenced by *SMN2-FL* mRNA levels and treatment.

### Treatment responses of individual cell lines vary substantially

We next compared treatment response to *SMN2* splice-modifying drugs between each of the 35 SMA patient-derived primary fibroblast cell lines that we included in our study. There was no significant difference in SMN levels after *in vitro* treatment with either drug in 15 cell lines (43%, **Fig. 3A**). In contrast, 20 of the cell lines (57%) showed a preference towards one of the treatments, as illustrated by a statistically significant difference in SMN protein levels between samples obtained from the same patients but treated *in vitro* with a different *SMN2* splice-modifying drug (**Fig. 3B** and **3C**). The number of cell lines preferably responding to nusinersen (31%) and risdiplam (26%) was similar, and we measured a difference of 41% of SMN level on average amongst cell lines that showed a preferred response to either treatment. SMN levels could differ as much as 85% (e.g. SMA_15 for nusinersen) or 72% (e.g. SMA_5 for risdiplam) in the same cell line but treated with a different drug. In control cell lines, we observed similar variability, as 60% showed a preference to one of the treatments, as illustrated by a statistically significant difference in SMN levels after nusinersen or risdiplam treatment (**Fig. S7**). We found no statistically significant enrichment for age, sex, SMA type or *SMN2* copy number in each of these groups. Interestingly, 8 of our 35 cell lines included a *SMN1-SMN2* hybrid gene copy, 7 of which showed a preference for either nusinersen or risdiplam. Our understanding of the clinical relevance of such hybrid genes remains limited, but this observation may provide an interesting starting point for further studies.

**Figure 3.**
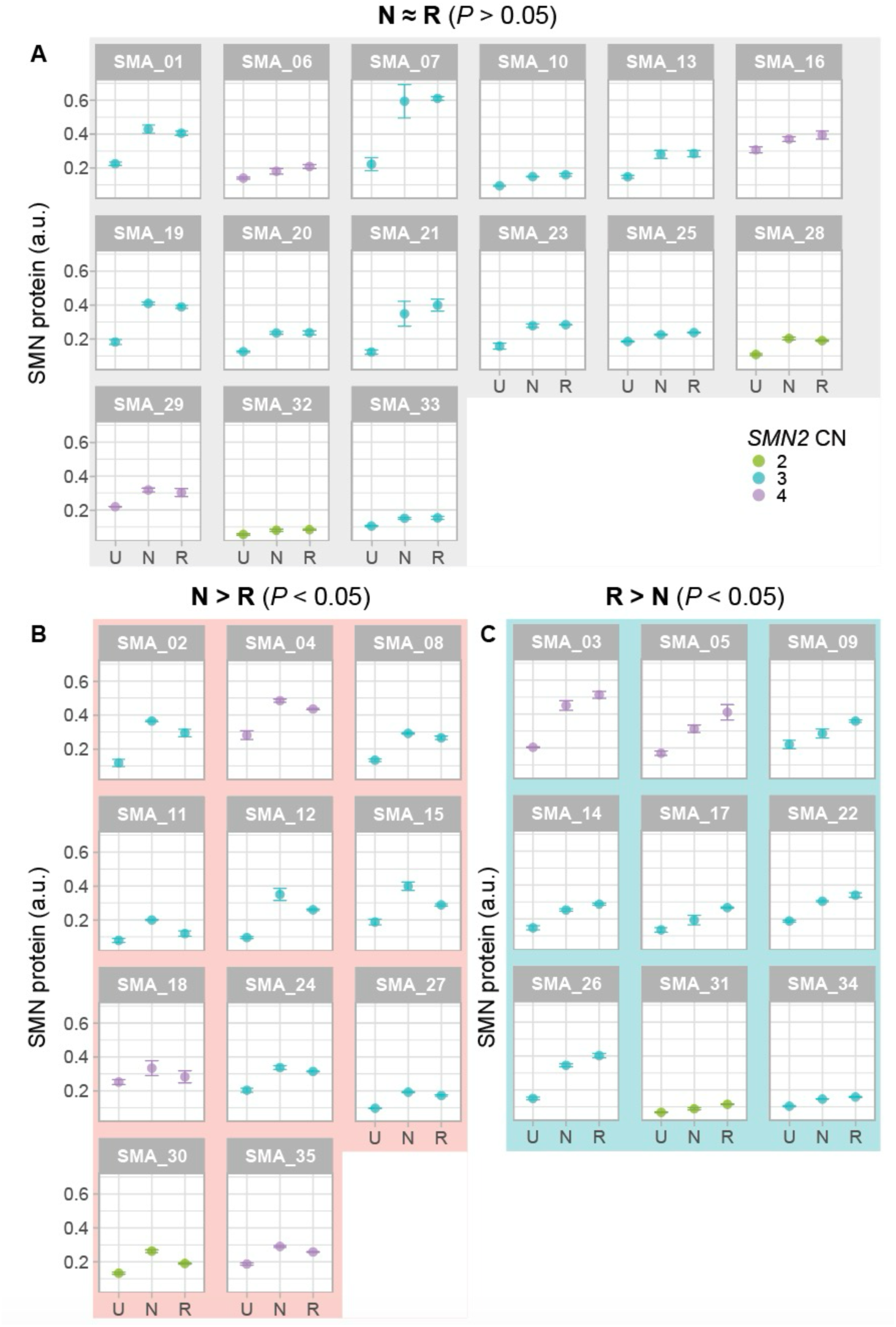
Variable responses to *SMN2* splice-modifying treatments in individual cell lines. **(A)** Cell lines for which *in vitro* treatment response increased SMN expression significantly compared to untreated condition, but with no significant difference between both treatments (15 cell lines, 43%). **(B)** Cell lines for which *in vitro* treatment response increased SMN expression significantly compared to untreated condition, and for which nusinersen treatment increased SMN expression more than risdiplam treatment (*P* < 0.05, 11 cell lines, 31%). **(C)** Cell lines for which *in vitro* treatment response increased SMN expression significantly compared to untreated condition, and for which risdiplam treatment increased SMN expression more than nusinersen treatment (*P* < 0.05, 9 cell lines, 26%). Data are represented as the average of the technical triplicate ± standard deviation. U = untreated; N = nusinersen; R = risdiplam; a.u. = arbitrary unit; CN = copy number.

### Younger age, higher SMN2 copy number and higher pre-treatment SMN protein levels predict in vitro treatment response

Lastly, we explored what other molecular and clinical characteristics of the patients included in our study would add to our understanding of treatment variability. First, we noticed that SMA fibroblasts with three *SMN2* copies demonstrated a more variable, but also a more pronounced relative change in SMN protein after treatment compared to SMA fibroblasts with two or four *SMN2* copies (**Fig. 4A**). In line with this, we observed that patients with SMA type 2 showed a higher relative change in SMN protein post-treatment than patients with more (type 1b) or less (type 4) severe SMA (**Fig. 4B**). Furthermore, we compared treatment response as expressed by SMN relative change to age, sex or the presence of hybrid genes (**Fig. 4C-E**) but did not identify any statistically significant correlations. To investigate if *in vitro* treatment response could predict treatment responses observed in patients *in vivo*, we correlated the increase in motor scores of children on nusinersen treatment (Scheijmans *et al*, 2022) with SMN relative change in the corresponding fibroblast cell lines following *in vitro* nusinersen treatment (**Fig. 4F**). Although this highlighted the challenges associated with this type of analysis – e.g. a limited number of patients, varying motor scales and variable treatment duration will all need to be considered – we found a possible correlation between increase in a common functional motor scale for SMA (Hammersmith functional motor scale enhanced, HFMSE) and SMN relative increase in this subgroup of patients (R^2^ = 0.42, *P* = 0.058), suggesting that *in vitro* analysis may indeed reflect *in vivo* outcomes.

**Figure 4.**
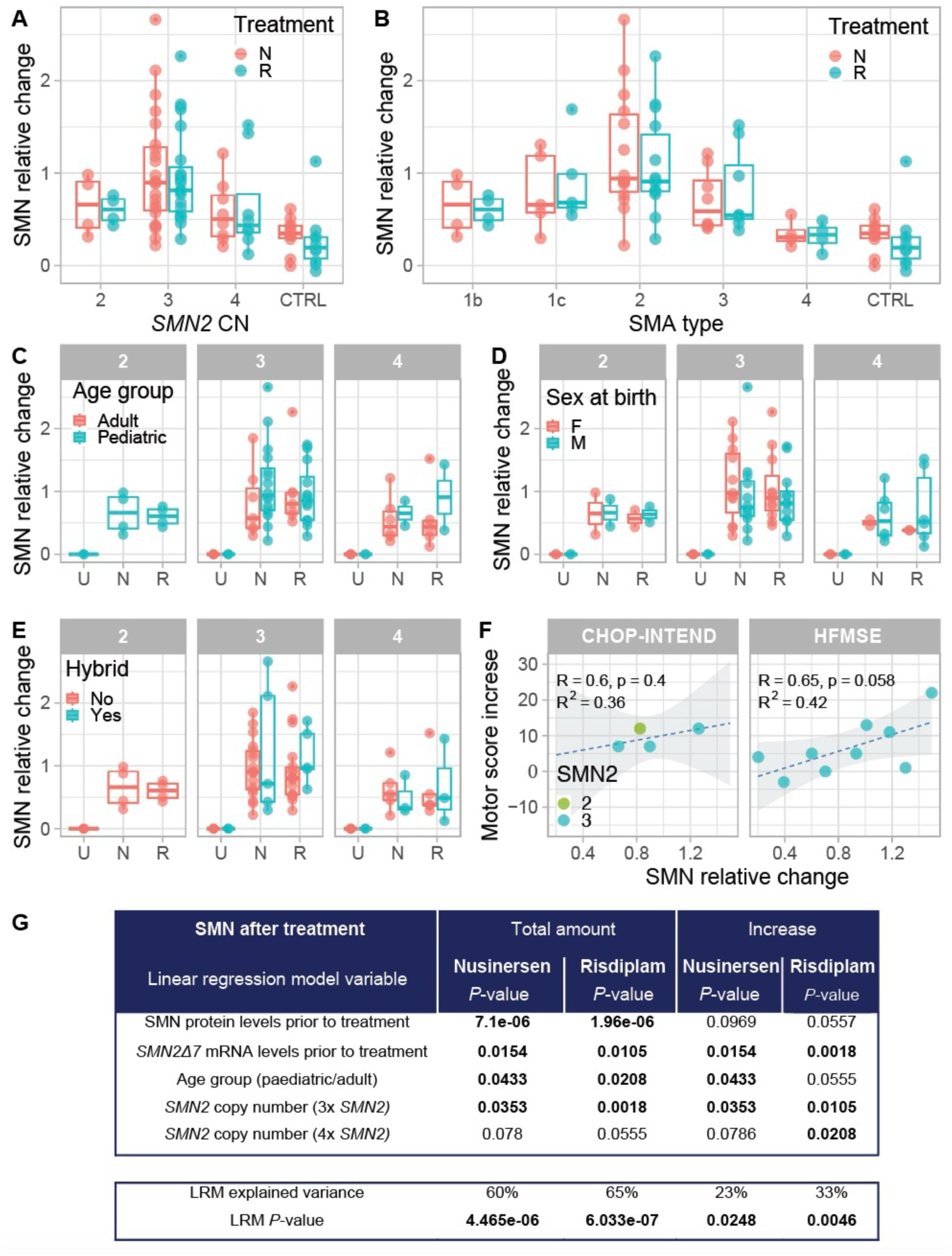
Identification of molecular factors influencing treatment effect. **(A)** SMN relative change in control (n=10) and SMA patient-derived primary fibroblasts with 2 *SMN2* copies (n=4), 3 *SMN2* copies (n=23) and 4 *SMN2* copies (n=8) after *in vitro* treatment with nusinersen (N, coral) or risdiplam (R, light blue). Each dot corresponds to the average of technical triplicate of each cell line. **(B)** SMN protein increase (relative change) in control (n=10) and SMA type 1b (n=4), type 1c (n=5), type 2 (n=14), type 3 (n=8) and type 4 (n=4) patient-derived primary fibroblasts after *in vitro* treatment with nusinersen (N) or risdiplam (R). Each dot corresponds to the average of the technical triplicate of each cell line. **(C)** SMN protein increase (relative change) in SMA patient-derived primary fibroblasts with 2x *SMN2* copies (n=4), 3x *SMN2* copies (n=23) and 4x *SMN2* copies (n=8) obtained from pediatric (light blue) and adult (coral) patients before treatment (U) and after *in vitro* treatment with nusinersen (N) and risdiplam (R). There is no statistically significant difference between pediatric and adult patients (one-way ANOVA, 3x*SMN2* nusinersen *P* = 0.28, risdiplam *P* = 0.92; 4x*SMN2* nusinersen *P* = 0.75, risdiplam *P* = 0.46). **(C)** SMN protein increase (relative change) in SMA patient-derived primary fibroblasts with 2x *SMN2* copies (n=4), 3x *SMN2* copies (n=23) and 4x *SMN2* copies (n=8) obtained from male (light blue) and female (coral) patients before treatment (U) and after *in vitro* treatment with nusinersen (N) and risdiplam (R). There is no statistically significant difference between pediatric and adult patients (one-way ANOVA, 2x*SMN2* nusinersen *P* = 0.99, risdiplam *P* = 0.74; 3x*SMN2* nusinersen *P* = 0.49, risdiplam *P* = 0.44; 4x*SMN2* nusinersen *P* = 0.75, risdiplam *P* = 0.46). **(E)** SMN protein increase (relative change) in SMA patient-derived primary fibroblasts with 2x *SMN2* copies (n=4), 3x *SMN2* copies (n=23) and 4x *SMN2* copies (n=8) with (light blue) and without (coral) *SMN2* hybrid genes before treatment (U) and after *in vitro* treatment with nusinersen (N) and risdiplam (R). There is no statistically significant difference between pediatric and adult patients (one-way ANOVA, 3x*SMN2* nusinersen *P* = 0.33, risdiplam *P* = 0.75; 4x *SMN2* nusinersen *P* = 0.61, risdiplam *P* = 0.35). **(F)** Correlation between motor score increase in patients after treatment with nusinersen and SMN protein increase (relative change). Data are represented as the average of the technical triplicate. Regression line (dashed line), Pearson correlation coefficient (R), p-value (p) and the coefficient of determination (R²) are displayed. (**G**) Linear regression model for SMN levels (total amount and increase) after *in vitro* treatment. *P* value of the factors significant in the linear regression model for SMN protein increase. *P* < 0.05 are highlighted in bold. U = untreated; N = nusinersen; R = risdiplam; a.u. = arbitrary unit. CHOP-INTEND = Children’s hospital of Philadelphia infant test of neuromuscular disorders. HFSME = Expanded Hammersmith Functional Motor Scale.

Finally, we investigated what molecular and clinical factors (age at biopsy, *SMN2* copy number, presence of *SMN1-SMN2* hybrid genes, sex, age, SMN levels prior to treatment, *SMN2* levels prior to treatment, *SMN2Δ7* levels prior to treatment, *SMN2* levels after treatment) influenced SMN protein levels after treatment (total protein levels and increase). We used linear regression models and used the backwise step approach to first systematically eliminate factors that played no statistically significant role (**Fig. 4G** and **Table S3**). We found that SMN protein and *SMN2Δ7* mRNA levels before treatment, age and *SMN2* copy number were statistically significant factors influencing levels of SMN protein after treatment. The interaction between these factors allows us to explain a limited percentage of variation of the increase of SMN levels after treatment (nusinersen: 23%, risdiplam: 33%). However, the interaction of SMN protein and *SMN2Δ7* mRNA before treatment, age and *SMN2* copy number predicted up to 60% (nusinersen, *P* = 4.47e-06) or 65% (risdiplam, *P* = 6.03e-07) of total SMN levels post-treatment. Like the correlation between total SMN levels and other factors (**Fig. S6**), these results point towards patient-specific factors regulating overall SMN protein expression that are not directly influenced by treatment.

## Discussion

In this study, we comprehensively characterized a large cohort of SMA patient- and healthy donor-derived fibroblasts following *in vitro* treatment with *SMN2* splice-modifying treatments. After treatment, *SMN2-FL* mRNA and SMN protein levels were significantly more heterogeneous than before treatment, reflecting variable outcomes in the SMA patient population. Importantly, more than half of the cell lines included in our study showed a preferred response to one of the two treatments examined, despite their similar mode of action. Most of the variation in levels of SMN protein after treatment was explained by younger age, higher *SMN2* copy number and higher pre-treatment SMN protein levels. We believe our results illustrate the importance of studying the effect of gene-targeting therapies in relevant model systems to enhance our understanding of fundamental molecular characteristics of SMA and how these are influenced by current gene-targeting therapies, aiming to identify objective measures of treatment outcomes that can support clinical decision making.

Our data highlights the potential of patient-derived fibroblasts for studying SMA and treatment responses when used in a sufficiently large number of different cell lines to account for common variation that occurs in the patient population. Indeed, previous studies have often been limited by their use of a small number of cell lines sourced primarily from repositories (Adami & Bottai, 2019; James *et al*, 2024; Kordala *et al*, 2023; Ottesen *et al*, 2023; McCormack *et al*, 2021), which may lack detailed clinical data about the donors and may not always express mRNA and protein levels representative for their genotype. To our knowledge, the number of studies including more than five patient-derived cell lines is limited (e.g. (Brown *et al*, 2022; Garbes *et al*, 2013; Wadman *et al*, 2016)). Garbes *et al*. found a correlation between SMA patient-derived fibroblasts and patient responses to valproic acid (VPA) treatment, highlighting the potential of using patient-derived fibroblasts to capture diverse treatment outcomes in SMA (Garbes *et al*, 2013). More recently, when using a large cohort of patient-derived fibroblasts to determine proteomic changes in SMA from different repositories and biobanks, Brown *et al*. found that SMN protein levels – and associated proteomic profiles – varied considerably between patients of the same SMA type and genotype (Brown *et al*, 2022). Our current dataset provides baseline reference data for future studies aimed at determining whether SMN expression of specific cell lines is representative for patients with a specific SMA type or *SMN2* copy number.

At baseline, we observed a limited correlation between *SMN2* mRNA and SMN protein levels. This finding aligns with previous observations, including for example in human *postmortem* spinal cord (Ramos *et al*, 2019), suggesting that factors beyond mRNA transcription and exon 7 splicing regulate SMN expression. Possible contributing factors may include the splicing of other *SMN2* exons, such as exon 3 and exon 5 (Ottesen *et al*, 2024), or involve translational (Ottesen *et al*, 2024; Workman *et al*, 2015) and post-translational regulation of the SMN protein (Rademacher *et al*, 2020; Detering *et al*, 2022). In line with this, we observed that *SMN2* copy number and SMA type did not linearly correlate with SMN protein increase after *in vitro* treatment. Rather, we noticed a higher SMN increase in cells with three *SMN2* copies or SMA type 2 than cells with two or four *SMN2* copies, and SMA type 1 or type 4. This may imply the presence of a molecular feedback loop that regulates SMN levels (Ottesen *et al*, 2024) and suggests that in addition to a minimum level of SMN that cells require for survival, there may also be a physiological maximum level of SMN that is deleterious to exceed. Indeed, neuronal toxicity has been reported in mouse studies of continuous, long-term AAV9-induced SMN overexpression (Van Alstyne *et al*, 2021; Xie *et al*, 2024; Zwartkruis & Groen, 2024). Studies to gain a better understanding of the regulation of SMN expression are vital, as they will have important implications for the optimisation of current gene-targeting therapies, and to identify novel therapeutic targets for the development of second-generation therapies for SMA.

Our ability to explain treatment outcomes using patient and clinical characteristics was incomplete. We found a possible link between sex and the regulation of SMN expression, through an enhanced correlation between *SMN2-FL* mRNA and SMN protein levels in male patients. This observation is interesting because of the role of key SMA modifiers UBA1 (Powis *et al*, 2016; Wishart *et al*, 2014) and plastin3 (Oprea *et al*, 2008; Hosseinibarkooie *et al*, 2016), located on the X chromosome. We are not aware of current studies reporting variable effects of gene-targeting treatment in patients of different sex, but our observation suggests this may warrant further analysis. Furthermore, we noticed a relationship between SMN levels and age. Interestingly, we found that in adults, SMN protein levels seemed relatively stable, as we and others reported before (Wadman *et al*, 2016). In children, however, we observed an initial increase in SMN expression, which is difficult to compare to previous studies as they been performed on blood samples (e.g. PBMC samples, e.g. (Crawford *et al*, 2012; Zaworski *et al*, 2016)). Our current data suggest a temporary increase in SMN levels between 2-14 years, which may follow on from a very high pre- and early post-natal requirement for SMN that was previously found to be followed by an immediate strong decrease in SMN levels in mostly young children (Ramos *et al*, 2019). Further refinement of the association between SMN and age will be important to better understand the uses and limitations of increasing SMN through gene-targeting therapies at later ages. Overall, it is likely that, despite including a relatively large number of cell lines in our experiments, our analyses remain underpowered with respect to detecting potential associations between molecular readouts and more subtle differences in patient characteristics. We believe, however, that the standardized cell culture approach we describe in this paper allows for the collection of samples over time, leading to the generation of such increasingly large datasets that will be required for refined and better-powered analyses in the future.

Patient-derived fibroblasts offer a relatively simple and scalable method to obtain primary cell lines whilst preserving genetic and epigenetic signatures of the donor (Franzen *et al*, 2021; Sturm *et al*, 2019), enabling personalized studies across many patients. The non-immortalized nature of primary fibroblasts leads to limitations on their growth potential, restricting expansion to a limited number of passages (Franzen *et al*, 2021), which needs to be monitored when working with large numbers of cell lines. Even though SMN is ubiquitously expressed and SMA is a systemic disease, motor neurons are the most affected cells (Hamilton & Gillingwater, 2013). Induced pluripotent stem cells (iPSCs) and iPSC-derived motor neurons (iPSC-MN) are therefore commonly used to study SMA (Adami & Bottai, 2019; Varderidou-Minasian *et al*, 2021; Son *et al*, 2019; Khayrullina *et al*, 2020; Zeng *et al*, 2023). However, their scalability is limited and most studies to date have been conducted using a limited number of cell lines. Many molecular changes observed in SMA-derived iPSC-MN have also been observed in primary fibroblasts, including a reduced number of nuclear gems (Ebert *et al*, 2009; Nizzardo *et al*, 2015; Coovert *et al*, 1997; Skordis *et al*, 2003), regulation of SMN expression by the long non-coding RNA *SMN-AS1* (d’Ydewalle *et al*, 2017), altered unfolded protein response (D’Amico *et al*, 2022; Ng *et al*, 2015), reduced mitochondrial function (Xu *et al*, 2016; Brown *et al*, 2022; Zilio *et al*, 2022), changes in ubiquitin-associated pathways (Powis *et al*, 2016; Brown *et al*, 2022; Fuller *et al*, 2016) and apoptotic defects. This highlights the possibilities of using fibroblasts as a scalable tool for the discovery of disease- and treatment-related molecular mechanisms (Signoria *et al*, 2023; Brown *et al*, 2022; Sansa *et al*, 2021; Sareen *et al*, 2012). However, the combination of appropriate animal- and human-based models will likely remain a requirement.

We found extensive variation in SMN increase after treatment between cell lines, including different responses to the two treatments. This suggests that patients can respond differently to *SMN2* -modifying treatments, despite both drugs targeting related sequences in and around exon 7 of *SMN2* (Singh *et al*, 2020, 2017) via comparable mechanisms. As there have been no clinical trials that directly compared SMN-targeting treatments, treatment choice currently remains mostly pragmatic and based on country-specific reimbursements (Yeo *et al*, 2024). Although the minimal SMN expression differences required for clinically relevant changes in patients are unknown, the robust differences we observed in many of our cell lines warrant speculation around switching treatments for certain patients. In a subset of cell lines, we were able to establish a correlation between *in vitro* treatment data with treatment response of patients from which we obtained the cell lines. A promising example for further development of this personalized medicine approach comes from research into the common genetic disease cystic fibrosis (CF). In CF, organoids derived from rectal biopsies were found to be highly suitable for measuring the effect of therapies on CF caused by specific genetic variants, and this approach has now been implemented in routine clinical decision-making around the start of and choice for specific treatments (Poel *et al*, 2023; Dekkers *et al*, 2016). Our preliminary analyses suggest that in the future, similar *in vitro* analyses may be used to assist decision making around choice or continuation of treatment for SMA.

In summary, our findings highlight the potential of patient-derived fibroblasts to study *in vitro* treatment efficacy. Our experiments suggest many patients may benefit more from one specific *SMN2* splice-modifying treatment, emphasizing the need to continue research that aims to identify objective measures to assist decision making around the choice and continuation of treatment. Although we identified molecular and clinical factors that predicted treatment outcomes *in vitro*, our understanding of variation in SMN levels before and after treatment outcomes remains incomplete. This warrants further studies into the cellular and molecular mechanisms that are associated with the regulation of SMN expression, and the influence of gene-targeting treatments on these mechanisms.

## Supporting information

all supplemental information

## Acknowledgements

We would like to thank all patients and their families for participating in our research and our research support staff for providing vital logistical and practical assistance to carry out this project. Our work was supported by grants from the European Union’s Horizon 2020 Research and Innovation Program under the Marie Skłodowska-Curie grant (H2020 Marie Skłodowska-Curie Actions) agreement no. 956185 (SMABEYOND ITN, to THG, GV, WLvdP, EG), Prinses Beatrix Spierfonds (W.OB21-01 to EJNG, RW, WLvdP), Stichting Spieren voor Spieren (to WLvdP), the Wellcome Trust (Edinburgh Clinical Academic Track (ECAT) to HM-O and THG), the Medical Research Council (MRC Clinician Scientist Fellows / MNDA Lady Edith Wolfson Clinical Fellow to KF), the Caitro Foundation (to EP and GV), and the European Union within the MUR PNRR ‘National Center for Gene Therapy and Drugs based on RNA Technology’ (Project no. CN00000041 CN3 RNA to GV).

## Materials and methods

### Ethical approval

Patients included for analysis in this study participate in a prospective, population-based study on SMA in the Netherlands. This study was approved by the UMC Utrecht Medical Ethical Committee (no. 09– 307/NL29692.041.09). We obtained additional written and oral informed consent for skin biopsies from each adult patient and from both parents of each participating minor.

### Study population and genetics

Patient characteristics were collected during interviews with parents using standardized questionnaires and physical examination as part of our ongoing population-based study (Wadman *et al*, 2017; Wijngaarde *et al*, 2020b, 2020a). Patients were classified as SMA types 1–4 based on the highest achieved motor milestone, following the international SMA classification system with some relevant additions (Mercuri *et al*, 2012; Wadman *et al*, 2017; Wijngaarde *et al*, 2020b). For all the subjects included, *SMN1* and *SMN2* copy number were confirmed using the SALSA multiplex ligation-dependent probe amplification (MLPA) kit P021 version B1 (MRC Holland) according to the manufacturer’s protocol and Coffyalyser.Net software (MRC Holland).

We selected samples from a cross-section of patients with varying *SMN2* copy number (2x *SMN2*: n=4, 3x *SMN2*: n=23; 4x *SMN2*: n=8). Of these patients, 9 had *SMN1-SMN2* gene hybrids (3 *SMN2* copies with one hybrid: n=5; 4 *SMN2* copies with one hybrid: n=4) and one had *SMN1* exon 1-6 deletion as determined by MLPA. The cell lines were from patients with SMA type 1b (n=4), SMA type 1c (n=5), SMA type 2 (n=14), SMA type 3 (n=8) and SMA type 4 (n=4). In addition, we analyzed cell lines from 10 healthy donors. Healthy donors had varying *SMN1* (2x *SMN1*: n=8, 3x *SMN1*: n=1, 4x *SMN1*: n=1) and *SMN2 (*1x *SMN2*: n=5, 2x *SMN2*: n=5) copies. Two controls had one full *SMN2* copy and one copy lacking exon 7 and exon 8 (*SMN2Δ7-8*); these are indicated in the figures as having 1x *SMN2*. The following cell lines were obtained from the NIGMS Human Genetic Cell Repository at the Coriell Institute for Medical Research: GM00232, GM03813, GM09677, GM03814, GM05659, GM03651, GM01650. The copy numbers for these cell lines were as indicated in the corresponding figure.

### Culture of primary fibroblasts

Primary fibroblasts were obtained from explants of 3 mm dermal biopsies. After 4-6 weeks, fibroblast outgrowths from the explants were enzymatically passaged (Accutase, Sigma Aldrich, A6964). Fibroblasts were cultured in DMEM containing 4.5 g/L glucose, L-glutamine and pyruvate (Gibco, 41966-029) with 10% fetal bovine serum (Cytvia, SH30073.03) and 1% penicillin-streptomycin (Sigma Aldrich, 0000165820). All cell lines were monitored and negative for mycoplasma (Merk, MP0035). We noticed changes in cellular growth rate and morphology at higher passages (>P13) and all our analyses were therefore performed on cells collected with low passage numbers (between P4 and P7). To maintain low passage numbers, and to collect cells for DNA, RNA, protein, and morphology at the same passage number, we developed the following workflow: cells were cultured and passaged up to 4xT175 flasks and grown until 80% of confluency. Cells were collected, counted and seeded as follows: 200,000 cells per 10 cm dish for protein collection; 100,000 cells in each well of a 6-well culture plate well for RNA isolation; 5,000 cells in each well of a 24-well plate well for morphological analysis; 400,000 cells in a T175 culture flask for DNA isolation (MLPA and biobanking). After treatment, cells were pelleted by washing twice with 1x PBS (Gibco, 10010-015), dissociated using Accutase and centrifugated for 10 min at 1,000x rcf at room temperature (RT). The pellet was again washed with 1x PBS and centrifuged for 10 min at 1,000x rcf at RT before storage at −80°C. To determine the relation between cell number, confluency and SMN expression, cells were collected and counted after 3, 4 and 5 days in culture. Subsequently, they were pelleted as described previously, stored at −80°C and SMN expression was determined as described below.

### Treatment with SMN2 splice-modifying drugs

Each cell line was cultured in three different conditions: untreated, treated with nusinersen (Biogen) or treated with risdiplam (Sanbio, 29028-1). Treatment with nusinersen and risdiplam was performed at 75 to 80% confluency. For nusinersen treatment, Lipofectamine LTX Reagent with PLUS Reagent (Invitrogen, A12621) was used following manufacturer’s recommendations at a final concentration of 10 nM. A 2′-O-methoxyethyl-modified scrASO (AGTTAGATGCCTATTCCU) was designed using GeneScript (www.genscript.com) and used as lipofection control. For risdiplam treatment, cells were treated at a final concentration of 0.5 µM (Sivaramakrishnan *et al*, 2017) and the treatment was repeated after 24 hours. The total treatment time for both treatments was 48 hours. After that, cells from 10 cm dishes and 6 well plates were pelleted and stored at −80°C for later analysis.

### RNA quantification using droplet digital PCR (ddPCR)

RNA was isolated using the RNeasy mini kit (Qiagen, 74104) following the manufacturer’s recommendations. RNA concentration was determined using a spectrophotometer (Nanodrop 2000, Thermo Scientific). Potential DNA contamination was prevented by DNaseI treatment (Thermo Scientific EN0521) of total RNA. cDNA was synthesized from 100 ng of RNA using the High-capacity cDNA reverse transcription kit (Applied Biosystems, 4368814) according to the manufacturer’s instructions. Primers and probes used for quantification of *SMN1*, *SMN2*, *SMN2Δ7* and *TBP* (housekeeping for normalisation) were obtained from IDT or Thermo Scientific (sequences as published previously (Ramos *et al*, 2019; Wadman *et al*, 2016), see **Table S1**). Reactions of 22 µL contained: 1 µL cDNA, 0.05 µL SMN-specific probe (100 µM), 0.05 µL TBP probe (100 µM), 1 µL of forward and reverse SMN- and TBP-specific primers (10 µM), 11 µL of 2x ddPCR Supermix for probes (no dUTP, Bio-Rad 186-3024) and 5.9 µL of RNase/DNase free water. Droplets were prepared using a QX200 Automated droplet generator (Bio-Rad 1864101). PCR was performed using a Bio-Rad T100 thermal cycler (95°C for 10 min, followed by 40 cycles of 95°C for 30 sec and 61.1°C for 1 min; followed by 98 °C for 10 min; ramp rate 2 °C/sec). After amplification, the droplets were analyzed using a QX200 droplet reader (Bio-Rad 1864003). Expression level of each *SMN* product (*SMN1*, *SMN2*, *SMN2Δ7*) was normalized to *TBP* expression using QuantaSoft Software (Bio-Rad 1864011). All experiments were run in technical triplicates.

### Protein quantification

Semi-quantitative western blotting was performed as described before (Groen *et al*, 2018). Cell pellets from 10 cm dishes were thawed on ice and homogenized in RIPA buffer (Thermo Scientific, 89900) with 1x protease inhibitor (Thermo Scientific, 1861278). Following incubation on ice for 10 min, the samples were centrifuged for 10 min at 4°C at 18,620x rcf. The supernatants were collected, and protein concentration was determined using the micro BCA protein assay kit (Thermo Scientific, 23235) following manufacturer’s recommendations. Protein concentration was normalized for all samples at 1 µg/µL in MilliQ water with 1x Bolt LDS sample buffer (Invitrogen, B0007) containing 1:20 β-mercaptoethanol (Sigma-Aldrich, M3148) and samples were incubated at 70°C for 10 minutes. Next, 20 µg of protein was loaded onto a Bolt bis-tris plus mini protein 4-12% gradient gel (Invitrogen, NW04125BOX), and samples were size-separated by running for 27 min at 200V in Bolt MES-SDS running buffer (Invitrogen, B0002). Proteins were subsequently transferred to a transfer stack containing a PDVF membrane (Invitrogen, IB24001) using the iBlot 2 gel transfer device (Invitrogen IB21001). Immediately after transfer, PDVF membranes were incubated in Revert 700 total protein stain (Li-Cor, 926-11011) for 5 min at RT, washed twice with washing buffer (30% methanol and 6.7% glacial acetic acid) and blocked in Odyssey PBS blocking buffer 1:3 in 1x PBS (Li-Cor, 927-40000) or EveryBlot blocking buffer 1:3 in 1x PBS (Biorad, 12010020) for 30 min (Odyssey) or 5 min (EveryBlot) at RT. Membranes were imaged with the Odyssey M imaging system (Li-Cor). Next, membranes were incubated in freshly made SMN-antibody solution (mouse-anti-SMN, BD Bioscience 610647, 1:1000 in blocking buffer) and incubated overnight at 4°C on rotation. After primary antibody incubation, membranes were washed three times for 10 min in PBS at RT and incubated in donkey-anti-mouse IRDye 800 (Li-Cor, 926-32212) secondary antibody diluted 1:2500 in blocking buffer with 0.02% SDS for 1 hour at RT. Membranes were finally washed 3 times for 30 min in PBS at RT and imaged with the Odyssey M imaging system. All experiments were run in technical triplicates. To facilitate reliable comparison of quantifications obtained from different membranes, results were normalised to an internal standard (SMN levels from HEK293 cell lysates) that was the same across all membranes included in our analyses. The internal standard allows to correct for technical variation caused by transfer, handling and processing of the individual western blotting membranes. Briefly, total protein staining (TPS) and SMN intensity were first determined using ImageStudio v5.2 software (Li-Cor). Then, SMN levels were normalized to the TPS intensity to control for loading variation. After normalizing SMN expression levels using TPS intensity for all samples, including the internal standard, the average intensity value of the internal standard was calculated and defined as 1 on each of the membranes. The SMN levels were then divided by the average value of the internal standard on each of the membranes, thus allowing to compare SMN levels across membranes (Huang *et al*, 2019). These values (in arbitrary units, a.u.) were used in the further statistical analyses as described later. All uncropped western blots used for quantification are included in **Supplementary figure 8**.

SMN ELISA was performed using the standardized SMN ELISA kit (2012, #ADI-900-209, Enzo Life Sciences, Farmingdale, NY) following the manufacturer’s instructions (Wadman *et al*, 2016).

### Immunofluorescence and microscopy

For morphological analysis, fibroblasts were stained for F-actin and DAPI. First, when cells reached 30 to 40% confluency, cells were fixed with 4% paraformaldehyde (PFA, Elektron Microscopy Sciences, 15710) at RT for 15 minutes and washed three times with 1x PBS. Next, cells were permeabilized with 0.1% Triton-X100 (Riedel-de Haën, 56029) for 5 min at RT, washed three times with 1x PBS and blocked with 2.5% bovine serum albumin (BSA) (Sigma Aldrich, CAS: 9048-46-8) for 60 minutes at RT. After blocking, the coverslip was incubated with 1.5% BSA containing 1:50 phalloidin A488 (Invitrogen, A12379) for 60 minutes at RT. Coverslips were washed three times with 1x PBS and incubated in 1.5% blocking buffer containing 300 nM DAPI (Invitrogen, D3571) for 10 minutes. After three washes with 1x PBS, the coverslips were mounted on a microscope slide with Mowiol 4-88 mounting medium (Sigma Aldrich, 81381) with 2.5% DABCO (Sigma Aldrich, D27802) and dried overnight. Three coverslips per cell line were imaged on a Leica DM inverted epifluorescent microscope. The nuclear and soma area were measured using Fiji (ImageJ2) version 2.3.0/1.53q by thresholding (Huang method) followed by measurement of the area and perimeter of the nucleus and soma. For confluency measurements, pictures were taken across a 10 cm tissue culture dish with a phase-contrast microscope. Images were thresholded manually to ensure coverage of the entire cell surface. Subsequently, the percentage of the image area covered by cells was quantified using Fiji (ImageJ2) version 2.3.0/1.53q.

### Statistical analysis

Statistical analysis was performed in R version 4.2.2 (2022-10-31). All data were tested for normality using the Shapiro-Wilk test. If normality was established, the appropriate parametric test was performed. For paired analysis, a two-tailed Welch’s t-test was conducted. In case of multiple comparisons, one-way ANOVA test followed by Tukey post-hoc test was performed. For treatment outcomes, a Welch pairwise t-test with Bonferroni’s correction for multiple testing was performed. For correlation analysis, Pearson correlation coefficient (R), p-value (p) and the coefficient of determination (R²) were estimated. For linear regression model building, the backward stepwise approach was used. Check of linearity of the data, independence and constant variance of residues was performed after the model building.

## References

Adami R & Bottai D (2019) Spinal Muscular Atrophy Modeling and Treatment Advances by Induced Pluripotent Stem Cells Studies. Stem Cell Rev and Rep 15: 795–813

Baranello G, Darras BT, Day JW, Deconinck N, Klein A, Masson R, Mercuri E, Rose K, El-Khairi M, Gerber M, et al (2021) Risdiplam in Type 1 Spinal Muscular Atrophy. New England Journal of Medicine 384: 915–923

Brown SJ, Kline RA, Synowsky SA, Shirran SL, Holt I, Sillence KA, Claus P, Wirth B, Wishart TM & Fuller HR (2022) The Proteome Signatures of Fibroblasts from Patients with Severe, Intermediate and Mild Spinal Muscular Atrophy Show Limited Overlap. Cells 11: 2624

Calucho M, Bernal S, Alías L, March F, Venceslá A, Rodríguez-Álvarez FJ, Aller E, Fernández RM, Borrego S, Millán JM, et al (2018) Correlation between SMA type and SMN2 copy number revisited: An analysis of 625 unrelated Spanish patients and a compilation of 2834 reported cases. Neuromuscular Disorders 28: 208–215

Chaytow H, Faller KME, Huang Y-T & Gillingwater TH (2021) Spinal muscular atrophy: From approved therapies to future therapeutic targets for personalized medicine. Cell Reports Medicine 2: 100346

Coovert DD, Le TT, McAndrew PE, Strasswimmer J, Crawford TO, Mendell JR, Coulson SE, Androphy EJ, Prior TW & Burghes AHM (1997) The Survival Motor Neuron Protein in Spinal Muscular Atrophy. Human Molecular Genetics 6: 1205–1214

Coratti G, Cutrona C, Pera MC, Bovis F, Ponzano M, Chieppa F, Antonaci L, Sansone V, Finkel R, Pane M, et al (2021) Motor function in type 2 and 3 SMA patients treated with Nusinersen: a critical review and meta-analysis. Orphanet Journal of Rare Diseases 16: 430

Crawford TO, Paushkin SV, Kobayashi DT, Forrest SJ, Joyce CL, Finkel RS, Kaufmann P, Swoboda KJ, Tiziano D, Lomastro R, et al (2012) Evaluation of SMN Protein, Transcript, and Copy Number in the Biomarkers for Spinal Muscular Atrophy (BforSMA) Clinical Study. PLOS ONE 7: e33572

Crawford TO, Swoboda KJ, De Vivo DC, Bertini E, Hwu W-L, Finkel RS, Kirschner J, Kuntz NL, Nazario AN, Parsons JA, et al (2023) Continued benefit of nusinersen initiated in the presymptomatic stage of spinal muscular atrophy: 5-year update of the NURTURE study. Muscle & Nerve 68: 157–170

D’Amico D, Biondi O, Januel C, Bezier C, Sapaly D, Clerc Z, El Khoury M, Sundaram VK, Houdebine L, Josse T, et al (2022) Activating ATF6 in spinal muscular atrophy promotes SMN expression and motor neuron survival through the IRE1α-XBP1 pathway. Neuropathology and Applied Neurobiology 48: e12816

Dekkers JF, Berkers G, Kruisselbrink E, Vonk A, de Jonge HR, Janssens HM, Bronsveld I, van de Graaf EA, Nieuwenhuis EES, Houwen RHJ, et al (2016) Characterizing responses to CFTR-modulating drugs using rectal organoids derived from subjects with cystic fibrosis. Science Translational Medicine 8: 344ra84–344ra84

Detering NT, Schüning T, Hensel N & Claus P (2022) The phospho-landscape of the survival of motoneuron protein (SMN) protein: relevance for spinal muscular atrophy (SMA). Cell Mol Life Sci 79: 497

Ebert AD, Yu J, Rose FF, Mattis VB, Lorson CL, Thomson JA & Svendsen CN (2009) Induced pluripotent stem cells from a spinal muscular atrophy patient. Nature 457: 277–280

Finkel RS, Mercuri E, Darras BT, Connolly AM, Kuntz NL, Kirschner J, Chiriboga CA, Saito K, Servais L, Tizzano E, et al (2017) Nusinersen versus Sham Control in Infantile-Onset Spinal Muscular Atrophy. New England Journal of Medicine 377: 1723–1732

Franzen J, Georgomanolis T, Selich A, Kuo C-C, Stöger R, Brant L, Mulabdić MS, Fernandez-Rebollo E, Grezella C, Ostrowska A, et al (2021) DNA methylation changes during long-term in vitro cell culture are caused by epigenetic drift. Commun Biol 4: 598

Fuller HR, Mandefro B, Shirran SL, Gross AR, Kaus AS, Botting CH, Morris GE & Sareen D (2016) Spinal Muscular Atrophy Patient iPSC-Derived Motor Neurons Have Reduced Expression of Proteins Important in Neuronal Development. Front Cell Neurosci 9

Garbes L, Heesen L, Holker I, Bauer T, Schreml J, Zimmermann K, Thoenes M, Walter M, Dimos J, Peitz M, et al (2013) VPA response in SMA is suppressed by the fatty acid translocase CD36. Human Molecular Genetics 22: 398–407

Groen EJN, Perenthaler E, Courtney NL, Jordan CY, Shorrock HK, Van Der Hoorn D, Huang Y-T, Murray LM, Viero G & Gillingwater TH (2018) Temporal and tissue-specific variability of SMN protein levels in mouse models of spinal muscular atrophy. Human Molecular Genetics 27: 2851–2862

Hamilton G & Gillingwater TH (2013) Spinal muscular atrophy: going beyond the motor neuron. Trends in Molecular Medicine 19: 40–50

Hosseinibarkooie S, Peters M, Torres-Benito L, Rastetter RH, Hupperich K, Hoffmann A, Mendoza-Ferreira N, Kaczmarek A, Janzen E, Milbradt J, et al (2016) The Power of Human Protective Modifiers: PLS3 and CORO1C Unravel Impaired Endocytosis in Spinal Muscular Atrophy and Rescue SMA Phenotype. The American Journal of Human Genetics 99: 647–665

Hua Y, Vickers TA, Okunola HL, Bennett CF & Krainer AR (2008) Antisense Masking of an hnRNP A1/A2 Intronic Splicing Silencer Corrects SMN2 Splicing in Transgenic Mice. The American Journal of Human Genetics 82: 834–848

Huang Y-T, van der Hoorn D, Ledahawsky LM, Motyl AAL, Jordan CY, Gillingwater TH & Groen EJN (2019) Robust Comparison of Protein Levels Across Tissues and Throughout Development Using Standardized Quantitative Western Blotting. J Vis Exp

James R, Faller KME, Groen EJN, Wirth B & Gillingwater TH (2024) Altered mitochondrial function in fibroblast cell lines derived from disease carriers of spinal muscular atrophy. Commun Med 4: 1–6

Khayrullina G, Moritz KE, Schooley JF, Fatima N, Viollet C, McCormack NM, Smyth JT, Doughty ML, Dalgard CL, Flagg TP, et al (2020) SMN-deficiency disrupts SERCA2 expression and intracellular Ca2+ signaling in cardiomyocytes from SMA mice and patient-derived iPSCs. Skelet Muscle 10: 16

Kobayashi DT, Olson RJ, Sly L, Swanson CJ, Chung B, Naryshkin N, Narasimhan J, Bhattacharyya A, Mullenix M & Chen KS (2011) Utility of Survival Motor Neuron ELISA for Spinal Muscular Atrophy Clinical and Preclinical Analyses. PLoS ONE 6: e24269

Kokaliaris C, Evans R, Hawkins N, Mahajan A, Scott DA, Sutherland CS, Nam J & Sajeev G (2024) Long-Term Comparative Efficacy and Safety of Risdiplam and Nusinersen in Children with Type 1 Spinal Muscular Atrophy. Adv Ther 41: 2414–2434

Kordala AJ, Stoodley J, Ahlskog N, Hanifi M, Garcia Guerra A, Bhomra A, Lim WF, Murray LM, Talbot K, Hammond SM, et al (2023) PRMT inhibitor promotes SMN2 exon 7 inclusion and synergizes with nusinersen to rescue SMA mice. EMBO Molecular Medicine 15: e17683

Kumbier K, Roth M, Li Z, Lazzari-Dean J, Waters C, Hammerlindl S, Rinaldi C, Huang P, Korobeynikov VA, Phatnani H, et al (2024) Identifying FUS amyotrophic lateral sclerosis disease signatures in patient dermal fibroblasts. Developmental Cell 0

Lefebvre S, Bürglen L, Reboullet S, Clermont O, Burlet P, Viollet L, Benichou B, Cruaud C, Millasseau P, Zeviani M, et al (1995) Identification and characterization of a spinal muscular atrophy-determining gene. Cell 80: 155–165

Lorson CL, Hahnen E, Androphy EJ & Wirth B (1999) A single nucleotide in the SMN gene regulates splicing and is responsible for spinal muscular atrophy. Proc Natl Acad Sci U S A 96: 6307–6311

McCormack NM, Abera MB, Arnold ES, Gibbs RM, Martin SE, Buehler E, Chen Y-C, Chen L, Fischbeck KH & Burnett BG (2021) A high-throughput genome-wide RNAi screen identifies modifiers of survival motor neuron protein. Cell Rep 35: 109125

Mendell JR, Al-Zaidy S, Shell R, Arnold WD, Rodino-Klapac LR, Prior TW, Lowes L, Alfano L, Berry K, Church K, et al (2017) Single-Dose Gene-Replacement Therapy for Spinal Muscular Atrophy. New England Journal of Medicine 377: 1713–1722

Mercuri E, Bertini E & Iannaccone ST (2012) Childhood spinal muscular atrophy: controversies and challenges. Lancet Neurol 11: 443–452

Mercuri E, Darras BT, Chiriboga CA, Day JW, Campbell C, Connolly AM, Iannaccone ST, Kirschner J, Kuntz NL, Saito K, et al (2018) Nusinersen versus Sham Control in Later-Onset Spinal Muscular Atrophy. New England Journal of Medicine 378: 625–635

Mercuri E, Sumner CJ, Muntoni F, Darras BT & Finkel RS (2022) Spinal muscular atrophy. Nat Rev Dis Primers 8: 1–16

Monani UR, Lorson CL, Parsons DW, Prior TW, Androphy EJ, Burghes AHM & McPherson JD (1999) A single nucleotide difference that alters splicing patterns distinguishes the SMA gene SMN1 from the copy gene SMN2. Human Molecular Genetics 8: 1177–1183

Naryshkin NA, Weetall M, Dakka A, Narasimhan J, Zhao X, Feng Z, Ling KKY, Karp GM, Qi H, Woll MG, et al (2014) SMN2 splicing modifiers improve motor function and longevity in mice with spinal muscular atrophy. Science 345: 688–693

Ng S-Y, Soh BS, Rodriguez-Muela N, Hendrickson DG, Price F, Rinn JL & Rubin LL (2015) Genome-wide RNA-Seq of Human Motor Neurons Implicates Selective ER Stress Activation in Spinal Muscular Atrophy. Cell Stem Cell 17: 569–584

Nizzardo M, Simone C, Dametti S, Salani S, Ulzi G, Pagliarani S, Rizzo F, Frattini E, Pagani F, Bresolin N, et al (2015) Spinal muscular atrophy phenotype is ameliorated in human motor neurons by SMN increase via different novel RNA therapeutic approaches. Sci Rep 5: 11746

Oprea GE, Kröber S, McWhorter ML, Rossoll W, Müller S, Krawczak M, Bassell GJ, Beattie CE & Wirth B (2008) Plastin 3 Is a Protective Modifier of Autosomal Recessive Spinal Muscular Atrophy. Science 320: 524–527

Oskoui M, Day JW, Deconinck N, Mazzone ES, Nascimento A, Saito K, Vuillerot C, Baranello G, Goemans N, Kirschner J, et al (2023) Two-year efficacy and safety of risdiplam in patients with type 2 or non-ambulant type 3 spinal muscular atrophy (SMA). J Neurol 270: 2531–2546

Ottesen EW, Seo J, Luo D, Singh NN & Singh RN (2024) A super minigene with a short promoter and truncated introns recapitulates essential features of transcription and splicing regulation of the SMN1 and SMN2 genes. Nucleic Acids Research 52: 3547–3571

Ottesen EW, Singh NN, Luo D, Kaas B, Gillette BJ, Seo J, Jorgensen HJ & Singh RN (2023) Diverse targets of SMN2-directed splicing-modulating small molecule therapeutics for spinal muscular atrophy. Nucleic Acids Research 51: 5948–5980

Poel E de, Spelier S, Hagemeijer MC, Mourik P van, Suen SWF, Vonk AM, Brunsveld JE, Ithakisiou GN, Kruisselbrink E, Oppelaar H, et al (2023) FDA-approved drug screening in patient-derived organoids demonstrates potential of drug repurposing for rare cystic fibrosis genotypes. Journal of Cystic Fibrosis 22: 548–559

Powis RA, Karyka E, Boyd P, Côme J, Jones RA, Zheng Y, Szunyogova E, Groen EJN, Hunter G, Thomson D, et al (2016) Systemic restoration of UBA1 ameliorates disease in spinal muscular atrophy. JCI Insight 1

Rademacher S, Detering NT, Schüning T, Lindner R, Santonicola P, Wefel I-M, Dehus J, Walter LM, Brinkmann H, Niewienda A, et al (2020) A Single Amino Acid Residue Regulates PTEN-Binding and Stability of the Spinal Muscular Atrophy Protein SMN. Cells 9: 2405

Ramos DM, d’Ydewalle C, Gabbeta V, Dakka A, Klein SK, Norris DA, Matson J, Taylor SJ, Zaworski PG, Prior TW, et al (2019) Age-dependent SMN expression in disease-relevant tissue and implications for SMA treatment. Journal of Clinical Investigation 129: 4817–4831

Sansa A, de la Fuente S, Comella JX, Garcera A & Soler RM (2021) Intracellular pathways involved in cell survival are deregulated in mouse and human spinal muscular atrophy motoneurons. Neurobiology of Disease 155: 105366

Sareen D, Ebert AD, Heins BM, McGivern JV, Ornelas L & Svendsen CN (2012) Inhibition of Apoptosis Blocks Human Motor Neuron Cell Death in a Stem Cell Model of Spinal Muscular Atrophy. PLOS ONE 7: e39113

Scheijmans FEV, Cuppen I, Van Eijk RPA, Wijngaarde CA, Schoenmakers MAGC, Van Der Woude DR, Bartels B, Veldhoen ES, Oude Lansink ILB, Groen EJN, et al (2022) Population-based assessment of nusinersen efficacy in children with spinal muscular atrophy: a 3-year follow-up study. Brain Communications 4: fcac269

Signoria I, van der Pol WL & Groen EJN (2023) Innovating spinal muscular atrophy models in the therapeutic era. Disease Models & Mechanisms 16: dmm050352

Singh NN, Howell MD, Androphy EJ & Singh RN (2017) How the discovery of ISS-N1 led to the first medical therapy for spinal muscular atrophy. Gene Ther 24: 520–526

Singh RN, Ottesen EW & Singh NN (2020) The First Orally Deliverable Small Molecule for the Treatment of Spinal Muscular Atrophy. J Exp Neurosci 15: 2633105520973985

Sivaramakrishnan M, McCarthy KD, Campagne S, Huber S, Meier S, Augustin A, Heckel T, Meistermann H, Hug MN, Birrer P, et al (2017) Binding to SMN2 pre-mRNA-protein complex elicits specificity for small molecule splicing modifiers. Nat Commun 8: 1476

Skordis LA, Dunckley MG, Yue B, Eperon IC & Muntoni F (2003) Bifunctional antisense oligonucleotides provide a trans-acting splicing enhancer that stimulates SMN2 gene expression in patient fibroblasts. Proceedings of the National Academy of Sciences 100: 4114–4119

Son YS, Choi K, Lee H, Kwon O, Jung KB, Cho S, Baek J, Son B, Kang S-M, Kang M, et al (2019) A SMN2 Splicing Modifier Rescues the Disease Phenotypes in an In Vitro Human Spinal Muscular Atrophy Model. Stem Cells Dev 28: 438–453

Strauss KA, Farrar MA, Muntoni F, Saito K, Mendell JR, Servais L, McMillan HJ, Finkel RS, Swoboda KJ, Kwon JM, et al (2022a) Onasemnogene abeparvovec for presymptomatic infants with two copies of SMN2 at risk for spinal muscular atrophy type 1: the Phase III SPR1NT trial. Nat Med 28: 1381–1389

Strauss KA, Farrar MA, Muntoni F, Saito K, Mendell JR, Servais L, McMillan HJ, Finkel RS, Swoboda KJ, Kwon JM, et al (2022b) Onasemnogene abeparvovec for presymptomatic infants with three copies of SMN2 at risk for spinal muscular atrophy: the Phase III SPR1NT trial. Nat Med 28: 1390–1397

Sturm G, Cardenas A, Bind M-A, Horvath S, Wang S, Wang Y, Hägg S, Hirano M & Picard M (2019) Human aging DNA methylation signatures are conserved but accelerated in cultured fibroblasts. Epigenetics 14: 961–976

Valori CF, Ning K, Wyles M, Mead RJ, Grierson AJ, Shaw PJ & Azzouz M (2010) Systemic Delivery of scAAV9 Expressing SMN Prolongs Survival in a Model of Spinal Muscular Atrophy. Sci Transl Med 2

Van Alstyne M, Tattoli I, Delestrée N, Recinos Y, Workman E, Shihabuddin LS, Zhang C, Mentis GZ & Pellizzoni L (2021) Gain of toxic function by long-term AAV9-mediated SMN overexpression in the sensorimotor circuit. Nat Neurosci 24: 930–940

Varderidou-Minasian S, Verheijen BM, Harschnitz O, Kling S, Karst H, Van Der Pol WL, Pasterkamp RJ & Altelaar M (2021) Spinal Muscular Atrophy Patient iPSC-Derived Motor Neurons Display Altered Proteomes at Early Stages of Differentiation. ACS Omega 6: 35375–35388

Wadman RI, Jansen MD, Stam M, Wijngaarde CA, Curial CAD, Medic J, Sodaar P, Schouten J, Vijzelaar R, Lemmink HH, et al (2020) Intragenic and structural variation in the SMN locus and clinical variability in spinal muscular atrophy. Brain Commun 2: fcaa075

Wadman RI, Stam M, Gijzen M, Lemmink HH, Snoeck IN, Wijngaarde CA, Braun KPJ, Schoenmakers MAGC, Van Den Berg LH, Dooijes D, et al (2017) Association of motor milestones, SMN2 copy and outcome in spinal muscular atrophy types 0–4. J Neurol Neurosurg Psychiatry 88: 365–367

Wadman RI, Stam M, Jansen MD, Van Der Weegen Y, Wijngaarde CA, Harschnitz O, Sodaar P, Braun KPJ, Dooijes D, Lemmink HH, et al (2016) A Comparative Study of SMN Protein and mRNA in Blood and Fibroblasts in Patients with Spinal Muscular Atrophy and Healthy Controls. PLoS ONE 11: e0167087

Wijngaarde CA, Stam M, Otto LAM, Bartels B, Asselman F-L, van Eijk RPA, van den Berg LH, Goedee HS, Wadman RI & van der Pol WL (2020a) Muscle strength and motor function in adolescents and adults with spinal muscular atrophy. Neurology 95: e1988–e1998

Wijngaarde CA, Stam M, Otto LAM, Van Eijk RPA, Cuppen I, Veldhoen ES, Van Den Berg LH, Wadman RI & Van Der Pol WL (2020b) Population-based analysis of survival in spinal muscular atrophy. Neurology 94: e1634–e1644

Wishart TM, Mutsaers CA, Riessland M, Reimer MM, Hunter G, Hannam ML, Eaton SL, Fuller HR, Roche SL, Somers E, et al (2014) Dysregulation of ubiquitin homeostasis and β-catenin signaling promote spinal muscular atrophy. J Clin Invest 124: 1821–1834

Workman E, Kalda C, Patel A & Battle DJ (2015) Gemin5 Binds to the Survival Motor Neuron mRNA to Regulate SMN Expression. J Biol Chem 290: 15662–15669

Xie Q, Chen X, Ma H, Zhu Y, Ma Y, Jalinous L, Cox GF, Weaver F, Yang J, Kennedy Z, et al (2024) Improved gene therapy for spinal muscular atrophy in mice using codon-optimized hSMN1 transgene and hSMN1 gene-derived promotor. EMBO Molecular Medicine 16: 945–965

Xu C-C, Denton KR, Wang Z-B, Zhang X & Li X-J (2016) Abnormal mitochondrial transport and morphology as early pathological changes in human models of spinal muscular atrophy. Disease Models & Mechanisms 9: 39–49

d’Ydewalle C, Ramos DM, Pyles NJ, Ng S-Y, Gorz M, Pilato CM, Ling K, Kong L, Ward AJ, Rubin LL, et al (2017) The Antisense Transcript SMN-AS1 Regulates SMN Expression and Is a Novel Therapeutic Target for Spinal Muscular Atrophy. Neuron 93: 66–79

Yeo CJJ, Tizzano EF & Darras BT (2024) Challenges and opportunities in spinal muscular atrophy therapeutics. The Lancet Neurology 23: 205–218

Zaworski P, Von Herrmann KM, Taylor S, Sunshine SS, McCarthy K, Risher N, Newcomb T, Weetall M, Prior TW, Swoboda KJ, et al (2016) SMN Protein Can Be Reliably Measured in Whole Blood with an Electrochemiluminescence (ECL) Immunoassay: Implications for Clinical Trials. PLoS ONE 11: e0150640

Zeng W, Kong X, Alamana C, Liu Y, Guzman J, Pang PD, Day JW & Wu JC (2023) Generation of two induced pluripotent stem cell lines from spinal muscular atrophy type 1 patients carrying no functional copies of SMN1 gene. Stem Cell Res 69: 103095

Zilio E, Piano V & Wirth B (2022) Mitochondrial Dysfunction in Spinal Muscular Atrophy. International Journal of Molecular Sciences 23: 10878

Zwartkruis MM & Groen EJ (2024) Promoting expression in gene therapy: more is not always better. EMBO Molecular Medicine 16: 672–674

