## Supplementary material for "Patient-specific responses to *SMN2* splice-modifying treatments in spinal muscular atrophy fibroblasts": all supplemental information

[1]Department of Neurology and Neurosurgery, UMC Utrecht Brain Center, and [2]Department of Genetics, University Medical Center Utrecht, Utrecht, the Netherlands; [3]Institute of Biophysics, CNR Unit, Trento, Italy; [4]Edinburgh Medical School: Biomedical Sciences and Euan MacDonald Centre for Motor Neurone Disease Research, and [5]Royal (Dick) School of Veterinary Studies, University of Edinburgh, Edinburgh, UK.

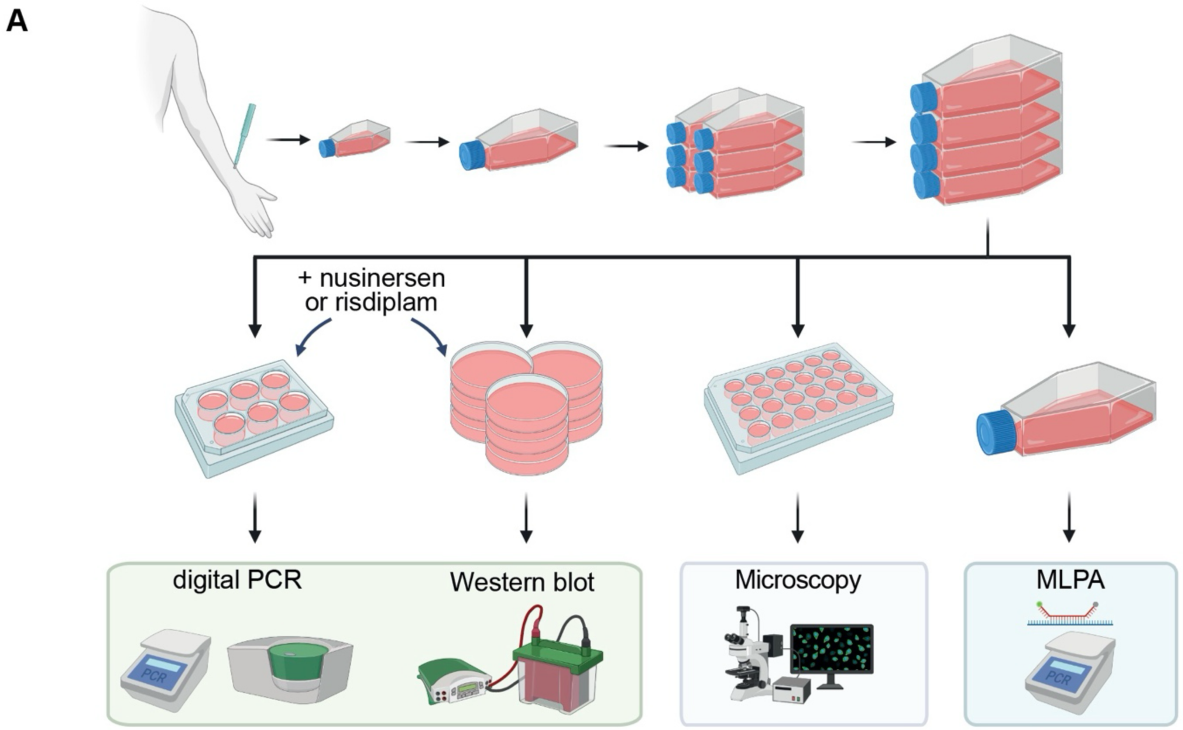


**Supplementary figure 1.** Workflow for the rapid expansion and culturing of primary-derived fibroblasts to obtain sufficient, low-passage numbers of cells as used in this study. Figure made with Biorender.com.

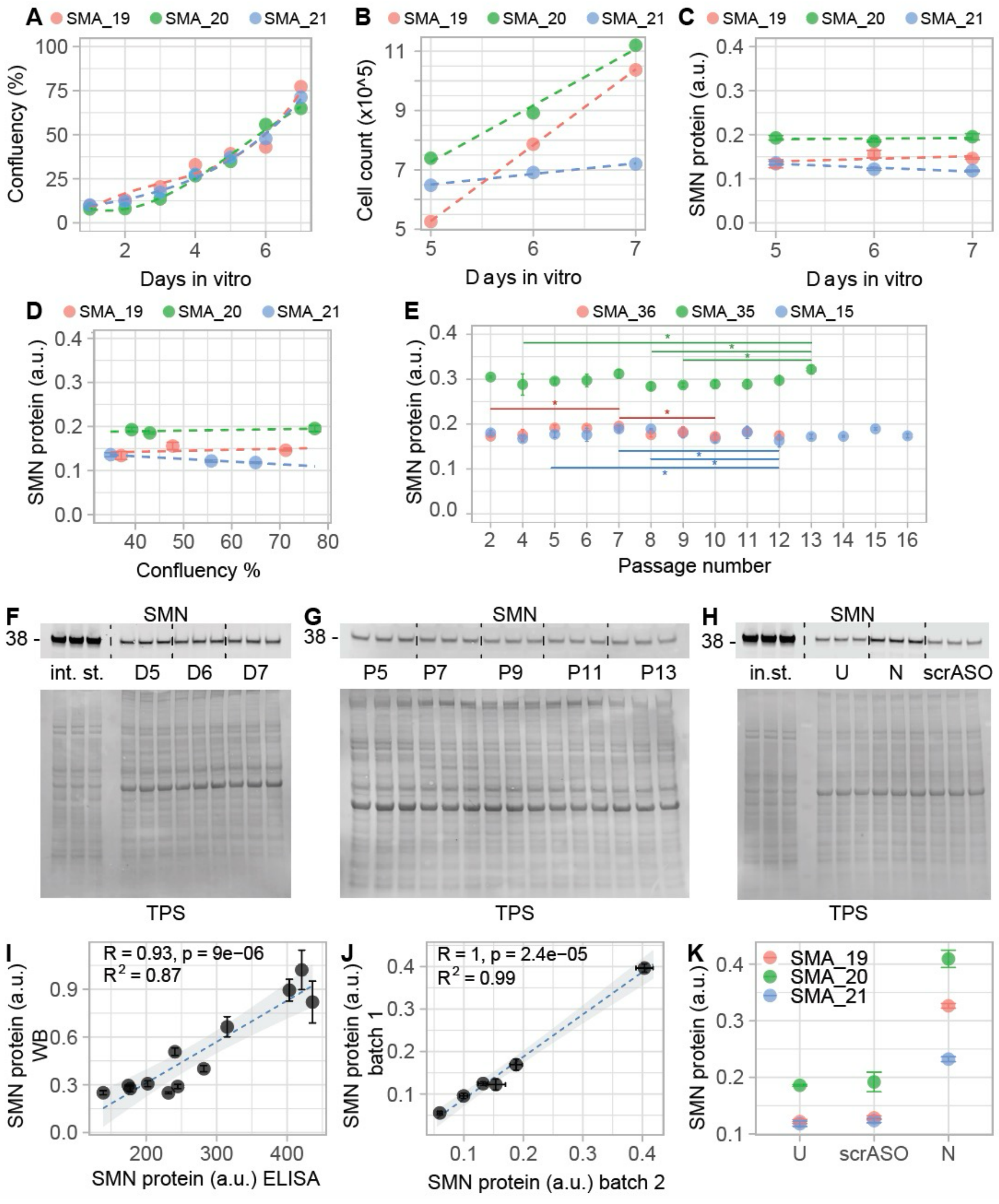


**Supplementary figure 2. Control analyses for SMN protein measurements in primary fibroblasts.** **(A)** Cell confluency measurement after 1 to 7 days *in vitro* of three different SMA patient-derived primary fibroblast lines. Confluency increases consistently over time in different cell lines. **(B)** Cell count over a 10cm dish after 5, 6 and 7 days *in vitro* of three different SMA patient-derived primary fibroblast lines. Cell number increase is variable, although within the same order of magnitude. Data are represented as the average of three biological triplicates. **(C)** Normalized SMN protein expression in SMA patient-derived primary fibroblast lines after 5, 6 and 7 days in culture. We found no statistically significant difference ($P$ = 0.43, two-way ANOVA). Data are represented as the average of technical triplicates ± standard deviation. **(D)** Relationship between normalized SMN protein expression in SMA patient-derived primary fibroblasts and cell confluency. SMN protein expression does not depend on confluency (*P* = 0.41, two-way ANOVA). Data are represented as the average of technical triplicates ± standard deviation. **(E)** Normalized SMN protein expression in SMA patient-derived primary fibroblasts at different passage numbers. We observed some variability but did not identify any consistent changes in SMN protein levels over time for the passage numbers we investigated (P2 to P16). Data are represented as the average of technical triplicates ± standard deviation. (Tukey post-hoc test: SMA_36 P7 vs. P2 *P* = 0.03, P10 vs. P7 *P* = 0.03; SMA_35 P12 vs. P7 *P* = 0.04, P12 vs. P8 *P* =0.04, P12 vs. P15 *P* = 0.04; SMA_15 P13 vs. P4 *P* = 0.05, P13 vs. P9 *P* = 0.04, P13 vs. P8 *P* = 0.02). **(F)** Representative western blot of SMN protein expression in SMA patient-derived primary fibroblasts at different days in vitro (D5, D6, D7) as used for analyses in (**C**) and (**D**). **(G)** Representative western blot of SMN protein expression in SMA patient-derived primary fibroblasts at different passage numbers (P5, P7, P9, P11, P13). **(H)** Representative western blot of SMN protein expression in SMA patient-derived primary fibroblast cell line untreated (U) and after *in vitro* treatment with either nusinersen (N) or a scrASO (scrASO). **(I)** Correlation between normalized SMN protein quantification via semi-quantitative western blotting (WB) and ELISA. The two methods show strong and significant correlation (*P* = 9e-06, $R^2$ = 0.87). Data are represented as the average of the technical triplicate ± standard deviation. The regression line (dashed line), its 95% confidence level interval, Pearson correlation coefficient (R), *P*-value (p) and the coefficient of determination ($R^2$) are displayed. **(J)** Correlation between SMN expression in two batches of the same cell lines thawed at two distinct time points. There is a strong and significant correlation (*P* = 2.4e-05, $R^2$ = 0.99). Data are represented as the average of the technical triplicate ± standard deviation. The regression line (dashed line), its 95% confidence level interval, Pearson correlation coefficient (R), *P*-value (p) and the coefficient of determination ($R^2$) are displayed. **(K)** SMN protein expression levels in three SMA patient-derived cell lines untreated (U), treated *in vitro* with either a scrASO or nusinersen (N) through lipofection. A representative western blot is shown in panel **H**. Data are represented as the average of the technical triplicate ± standard deviation. TPS = total protein staining; in. st = internal standard; N = nusinersen.; scrASO = scrambles antisense oligonucleotide; a.u. = arbitrary unit; * = *P* < 0.05.

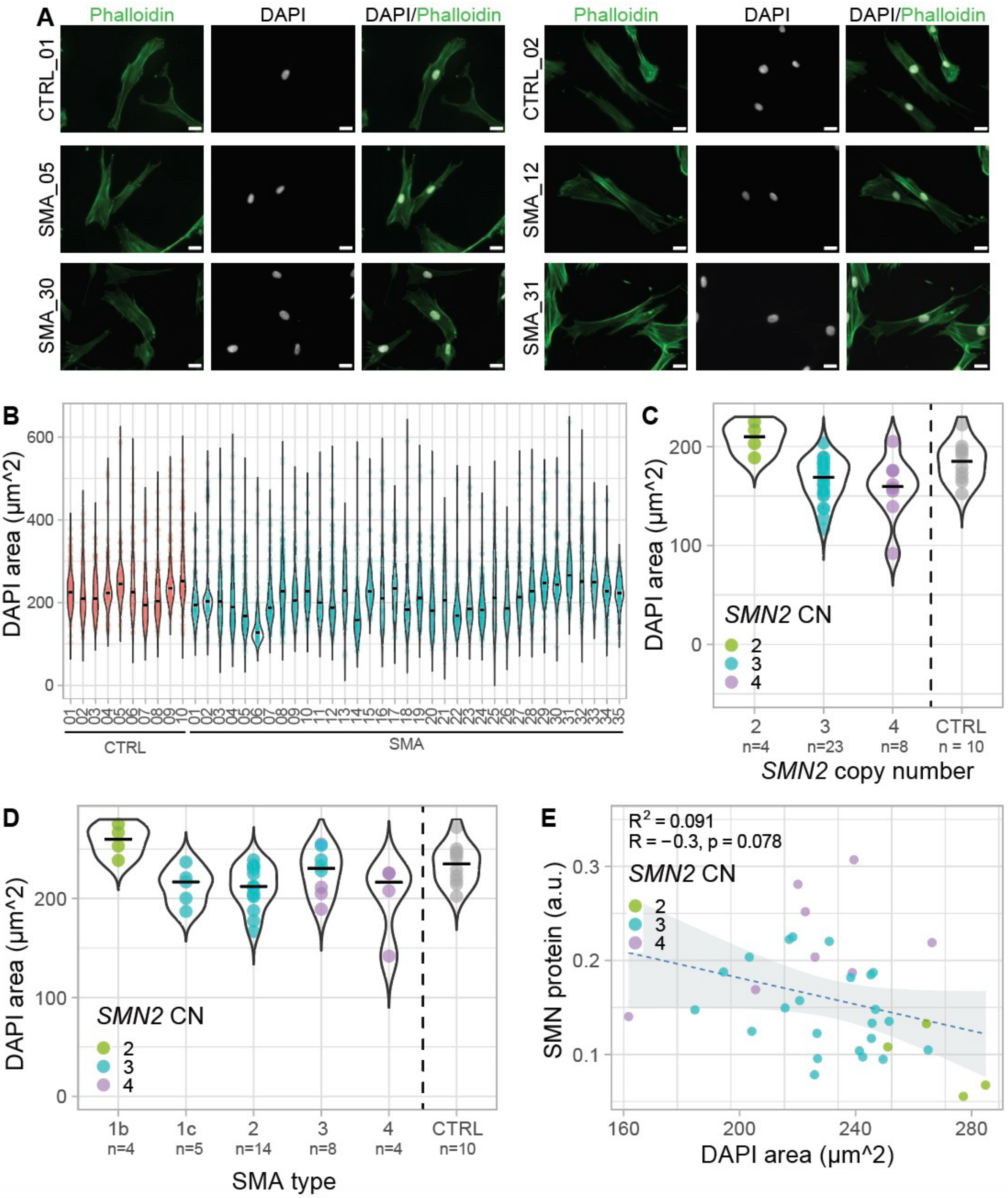


**Supplementary figure 3. Morphological characterization of control and SMA patient-derived fibroblasts. (A)** Representative images for control and SMA patient-derived fibroblasts. F-actin was labelled using phalloidin conjugated to Alexa Fluor 488. Chromatin was labelled using DAPI. SMA_05 and SMA_12 respectively have 4x *SMN2* and 3x *SMN2* copies. SMA_30 and SMA_31 have 2x *SMN2*. Scale bar: 25 µm. **(B)** DAPI (nucleus) area (µm²) in control and SMA patient-derived primary fibroblasts. Each dot corresponds to the area of one nucleus. **(C)** Average DAPI area (µm²) of control (n=10) and SMA patient-derived primary fibroblasts with 2x *SMN2* copies (n=4), 3x *SMN2* copies (n=23) and 4x *SMN2* copies (n=8). Nuclear size depends on *SMN2* copy number (one-way ANOVA, *P*=0.002, 2 vs. 3 p=0.009; 2 vs. 4 *P*=0.005). Each dot corresponds to the average nuclear size of each cell line. **(D)** DAPI area (μm²) of control (n=10) and SMA type 1b (n=4), type 1c (n=5), type 2 (n=14), type 3 (n=8) and type 4 (n=4) patient-derived primary fibroblasts. Nuclear size is partially related to SMA type (one-way ANOVA, *P*= 0.004, 1b vs. 1c *P*= 0.05; 1b vs. 2 *P*= 0.01; 1b vs. 4 p=0.01). Each dot corresponds to the average nuclear size of each cell line. **(E)** Relationship between normalized SMN protein expression levels (a.u.) and DAPI area (μm²) in SMA patient-derived primary fibroblasts. SMN protein does not correlate with nuclear area (*P* = 0.08). Data are represented as the average nuclear area of each cell line. The regression line (dashed line), its 95% confidence level interval, Pearson correlation coefficient (R), *P*-value (p) and the coefficient of determination ($R^2$) are displayed. CN = copy number.

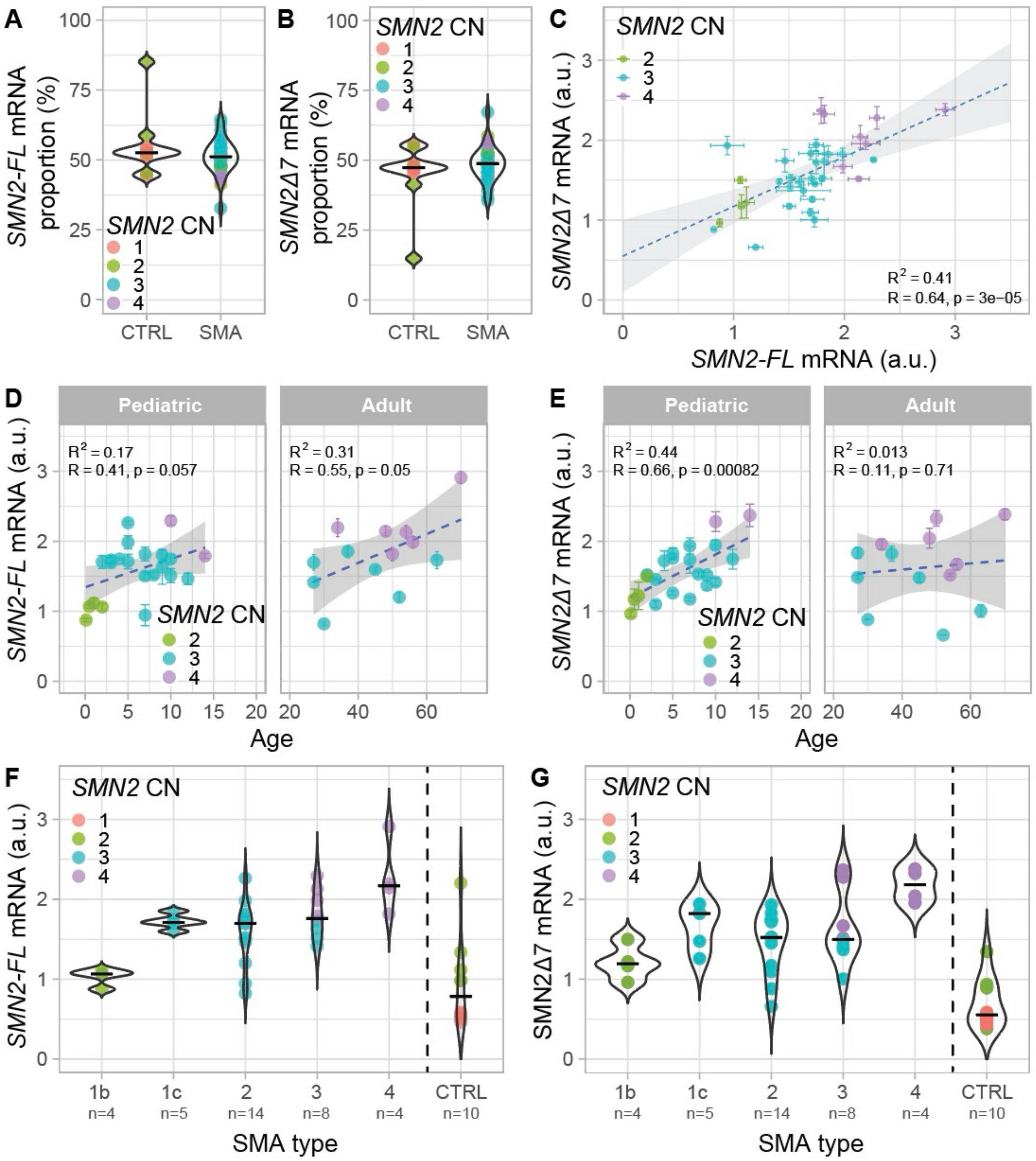


**Supplementary figure 4. *SMN2* mRNA expression in controls and SMA patient-derived primary fibroblasts. (A)** Percentage of *SMN2-FL* mRNA relative to total *SMN2* mRNA in control (n=10) and SMA patient-derived primary fibroblasts (n=35). There is no statistically significant difference in the relative abundance of *SMN2-FL* in control and SMA patient-derived fibroblasts. Each dot corresponds to the average of the technical triplicate of each cell line. **(B)** Percentage of *SMN2Δ7* mRNA relative to total *SMN2* mRNA in control (n=10) and SMA patient-derived primary fibroblasts (n=35). There is no statistically significant difference in the relative abundance of *SMN2Δ7* in control and patient-derived fibroblasts. Each dot corresponds to the average of technical triplicate of each cell line. **(C)** Relationship between *SMN2-FL* and *SMNΔ7* mRNA expression levels in SMA patient-derived primary fibroblasts. *SMN2-FL* and *SMN2Δ7* mRNA expression significantly correlates ($P$ = 3e-05, $R^2$ = 0.41). Data are represented as the average of the technical triplicate ± standard deviation. The regression line (dashed line), its 95% confidence level interval, Pearson correlation coefficient (R), *P*-value (p) and the coefficient of determination ($R^2$) are displayed. **(D)** Relationship between *SMN2-FL* mRNA expression and age at biopsy in SMA patient-derived primary fibroblasts. *SMN2-FL* mRNA expression shows trend of weak correlation with age in pediatric and adults ($P$ = 0.057 and $P$ = 0.05, respectively). Data are represented as the average of the technical triplicate ± standard deviation. Regression line (dashed line), Pearson correlation coefficient (R), p-value (p) and the coefficient of determination ($R^2$) are displayed. **(E)** Relationship between *SMN2Δ7* mRNA expression and age at the biopsy in SMA patient-derived primary fibroblasts. *SMN2Δ7* mRNA expression correlates with age in pediatric ($P$ = 0.0008) but not adult patients ($P$ = 0.71). Data are represented as the average of the technical triplicate ± standard deviation. The regression line (dashed line), Pearson correlation coefficient (R), *P*-value (p) and the coefficient of determination ($R^2$) are displayed. **(F)** *SMN2-FL* mRNA expression in control (n=10) and SMA type 1b (n=4), type 1c (n=5), type 2 (n=14), type 3 (n=8) and type 4 (n=4) patient-derived primary fibroblasts. There are differences in *SMN2-FL* mRNA expression across different SMA types (one-way ANOVA $P$ = 0.0003, 1b vs. 1c $P$ = 0.03, 1b vs. 2 $P$ = 0.045, 1b vs. 3 $P$ = 0.005, 1b vs. 4 $P$ = 0.0001, 2 vs. 4 $P$ = 0.00). Each dot corresponds to the average of technical triplicate of each cell line. **(G)** *SMN2Δ7* expression in control (n=10) and SMA type 1b (n=4), type 1c (n=5), type 2 (n=14), type 3 (n=8) and type 4 (n=4) patient-derived primary fibroblasts. There are some differences in *SMN2Δ7* mRNA expression across different SMA types (one-way ANOVA $P$ 0.009, 1b vs. 4 $P$ = 0.006, 2 vs. 4 $P$ = 0.01). Each dot corresponds to the average of technical triplicate of each cell line. a.u. = arbitry unit; CN = copy number.

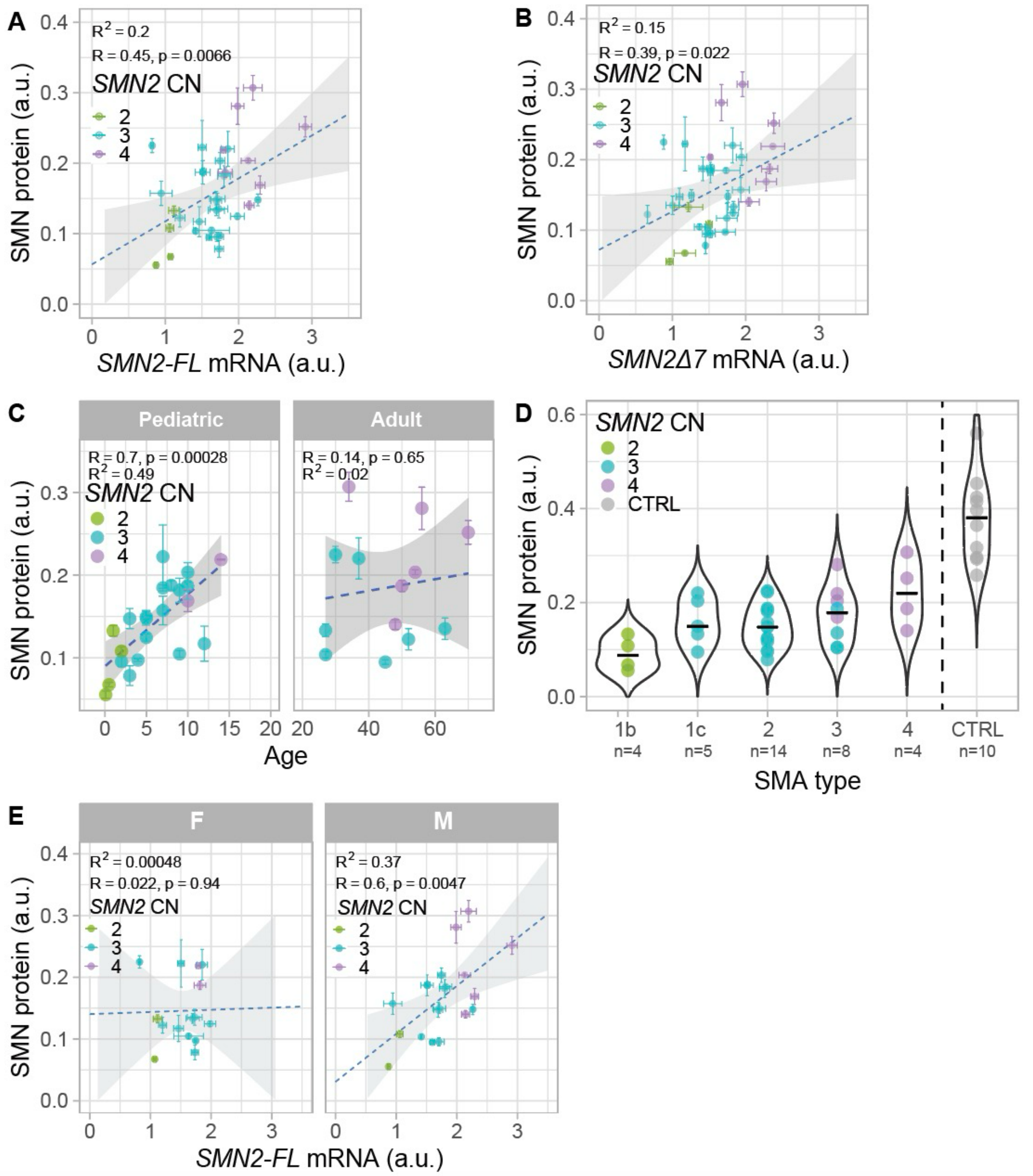


**Supplementary figure 5. SMN protein expression in controls and SMA patients-derived primary fibroblasts. (A)** Relationship between *SMN2-FL* mRNA and normalized SMN protein expression levels in SMA patient-derived primary fibroblasts. *SMN2-FL* mRNA significantly correlates with SMN protein levels (*P* = 0.007). Data are represented as the average of the technical triplicate ± standard deviation**.** The regression line (dashed line), its 95% confidence level interval, Pearson correlation coefficient (R), *P*-value (p) and the coefficient of determination (R²) are displayed. **(B)** Relationship between *SMN2Δ7* mRNA and normalized SMN protein expression levels in SMA patient-derived primary fibroblasts. *SMN2Δ7* mRNA correlates with SMN protein production (*P* = 0.02). Data are represented as the average of the technical triplicate ± standard deviation. The regression line (dashed line), its 95% confidence level interval, Pearson correlation coefficient (R), *P*-value (p) and the coefficient of determination ($R^2$) are displayed. **(C)** Correlation between normalized SMN protein expression and age at biopsy in SMA fibroblasts. Protein expression correlates with age in pediatric ($P$ = 0.003) but not adult patients ($P$ = 0.65). Data are represented as the average of the technical triplicate ± standard deviation. The regression line (dashed line), its 95% confidence level interval, Pearson correlation coefficient (R), p-value (p) and the coefficient of determination ($R^2$) are displayed. **(D)** Normalized SMN protein expression levels in control (n=10) and SMA type 1b (n=4), type 1c (n=5), type 2 (n=14), type 3 (n=8) and type 4 (n=4) patient-derived primary fibroblasts. SMN protein expression does not depend on SMA type (one-way ANOVA, $P$ = 0.02, 1b vs. 4 $P$ = 0.01). Each dot corresponds to the average of a technical triplicate for each cell line. **(E)** Relationship between *SMN2-FL* mRNA and normalized SMN protein expression levels in SMA patient-derived primary fibroblasts from female and male donors. *SMN2-FL* mRNA and SMN protein correlation is influenced by the donors' sex. Data are represented as the average of the technical triplicate ± standard deviation. The regression line (dashed line), its 95% confidence level interval, Pearson correlation coefficient (R), p-value (p) and the coefficient of determination ($R^2$) are displayed. TPS = total protein staining; a.u. = arbitrary unit; CN = copy number.

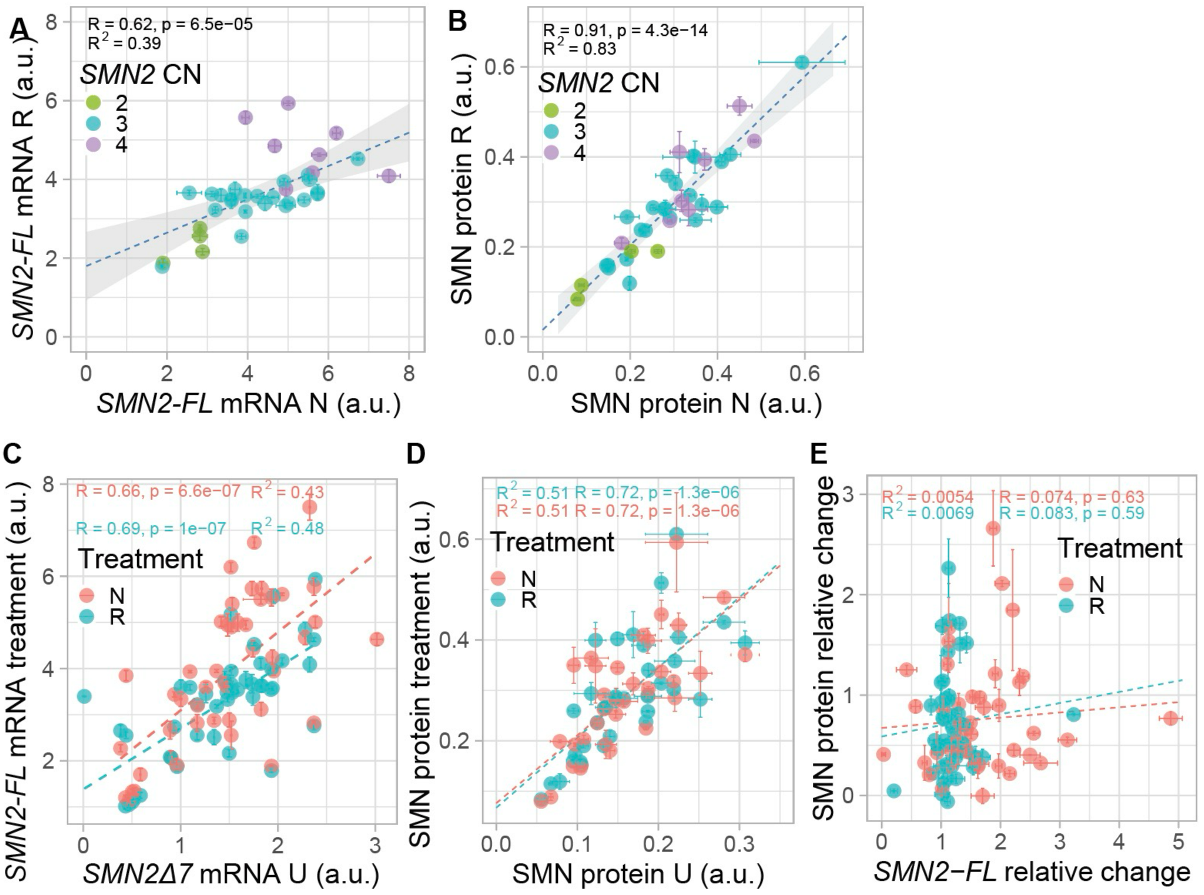

**Supplementary figure 6. The effect of *SMN2*-splice modifiers on *SMN2-FL*, *SMN2*Δ*7* and SMN expression levels in control and SMA patient-derived primary fibroblasts.**

**(A)** Relationship between *SMN2-FL* mRNA in SMA patient-derived primary fibroblasts treated with nusinersen (N) and risdiplam (R). Data are represented as the average of the technical triplicate ± standard deviation. The regression line, its 95% confidence level interval, Pearson correlation coefficient (R), p-value (p) and the coefficient of determination ($R^2$) are displayed. **(B)** Relationship between normalized SMN protein in SMA patient-derived primary fibroblasts treated with nusinersen (N) and risdiplam (R). Data are represented as the average of the technical triplicate ± standard deviation. Regression line, its 95% confidence level interval, Pearson correlation coefficient (R), *P*-value (p) and the coefficient of determination ($R^2$) are displayed. **(C)** Correlation between *SMN2-FL* mRNA after *in vitro* treatment with either nusinersen (N) and risdiplam (R) and *SMN*Δ*7* mRNA level in untreated fibroblasts (U) in SMA patient-derived fibroblasts. There is a statistically significant correlation. Data are represented as the average of the technical triplicate ± standard deviation. The regression line, Pearson correlation coefficient (R), *P*-value (p) and the coefficient of determination ($R^2$) are displayed. **(D)** Relationship between normalized SMN protein in untreated SMA patient-derived primary fibroblasts (U) and normalized SMN protein after *in vitro* treatment with risdiplam (R, light blue) or nusinersen (N, coral). Data are represented as the average of the technical triplicate ± standard deviation. Regression line, Pearson correlation coefficient (R), *P*-value (p) and the coefficient of determination ($R^2$) are displayed. **(E)** Relationship between *SMN2-FL* mRNA increase after *in vitro* treatment displayed as relative change and SMN protein increase after treatment displayed as relative change after treatment with either nusinersen (N, coral) and risdiplam (R, light blue) in SMA patient-derived primary fibroblasts. The correlation is not statistically significant for both treatments. Data are represented as the average of the technical triplicate ± standard deviation. Regression line, Pearson correlation coefficient (R), *P*-value (p) and the coefficient of determination ($R^2$) are displayed. U = untreated; N = nusinersen; R = risdiplam; a.u. = arbitrary unit; CN = copy number.

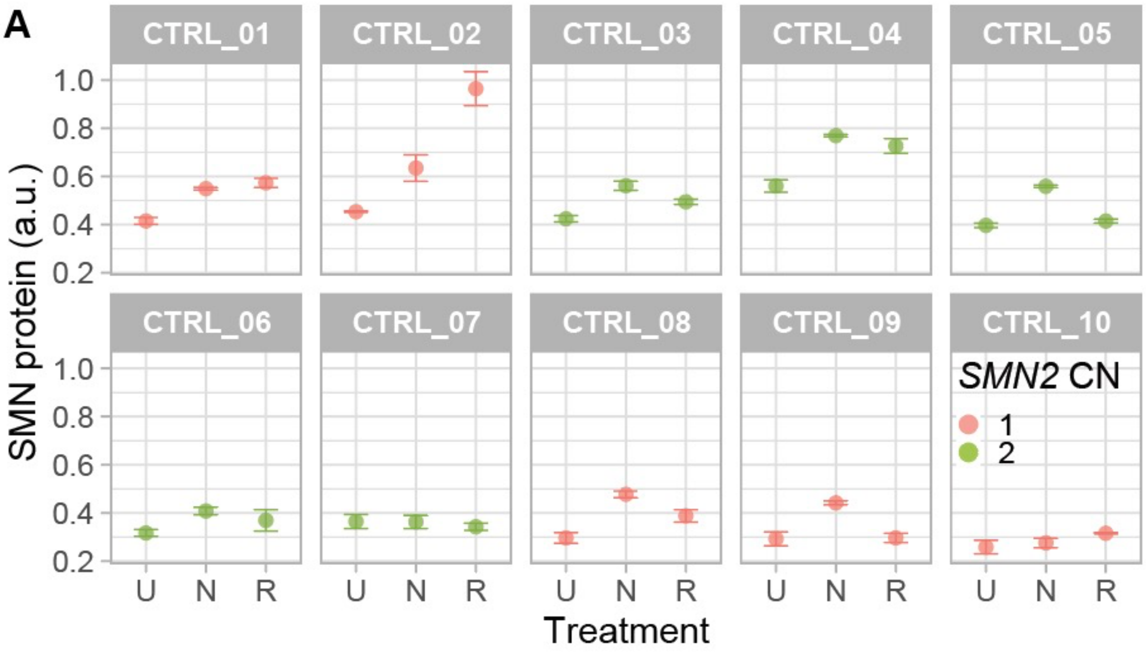


**Supplementary figure 7. Treatment effect across control cell lines.**

**(A)** Normalized SMN protein expression levels in control patient-derived primary fibroblasts untreated (U) and after *in vitro* treatment with either nusinersen (N) or risdiplam (R). Data are represented as the average of the technical triplicate ± standard deviation. U = untreated; N = nusinersen; R = risdiplam; a.u. = arbitrary unit; CN = copy number.

| Target | Primer/Prob | Sequence |
|---|---|---|
| *SMN1* | FW | TAC ATG AGT GGC TAT CAT ACT GGC TA |
| | Probe | 5'-6FAM TAT GGG TTT CAG ACA AA MGB-3' |
| | RV | AAT GTG AGC ACC TTC CTT CTT TTT |
| *SMN2* | FW | AC ATG AGT GGC TAT CAT ACT GGC TA |
| | Probe | 5'-6FAM ATA TGG GTT TTA GAC AAA A MGB-3' |
| | RV | AAT GTG AGC ACC TTC CTT CTT TTT |
| *SMNΔ7* | FW | GG CTA TCA TAC TGG CTA TTA TAT GGA A |
| | Probe | 5'-6FAM CTG GCA TAG AGC AGC ACT AAA TGA CAC CAC MGB-3' |
| | RV | TCC AGA TCT GTC TGA TCG TTT CTT |
| *TBP* | FW | CGT GGT TCG TGG CTC TCT |
| | Probe | 5’-HEX ATC CCA AGC -ZEN- GGT TTG CTG-3’ |
| | RV | GCC CGA AAC GCC GAA TAT |

**Table S1**. **Overview of primer and probe sequences used for ddPCR analyses.** Target name, primer or probe and sequences are indicated. FW = forward; RV = reverse. The probes are FAM-MGB (*SMN*) or HEX-ZEN (*TBP*). For references to previous studies using these sequences, please see the main text of our manuscript (**Materials and methods**).

| | | Type 1b (n=4) | Type 1c (n=5) | Type 2 (n=14) | Type 3 (n=8) | Type 4 (n=4) | Control (n=10) |
|---|---|---|---|---|---|---|---|
| Sex (M:F) | | 2:2 | 3:2 | 9:5 | 5:3 | 3:1 | 4:6 |
| *SMN1* CN | 0 | 4 | 5 | 14 | 8 | 4 | 0 |
| | 1 | 0 | 0 | 0 | 0 | 0 | 0 |
| | 2 | 0 | 0 | 0 | 0 | 0 | 8 |
| | 3 | 0 | 0 | 0 | 0 | 0 | 1 |
| | 4 | 0 | 0 | 0 | 0 | 0 | 1 |
| *SMN2* CN | 1 | 0 | 0 | 0 | 0 | 0 | 5 |
| | 2 | 4 | 0 | 0 | 0 | 0 | 5 |
| | 3 | 0 | 5 | 14 | 4 | 0 | 0 |
| | 4 | 0 | 0 | 0 | 4 | 4 | 0 |
| Median age in months at onset (range) | | 1.8 (0-2.5) | 6 (6-9) | 12 (6-36) | 39 (12-168) | 327 (246-510) | - |
| Median age in years at biopsy (range) | | 0.8 (0-2) | 27 (5-45) | 7 (2-32) | 20.5 (9-63) | 49 (34-70) | 28 (25-62) |

**Table S2. Overview of SMA and control clinical and genetic characteristics.**

| **SMN after treatment** | Total amount | | Increase | |
|---|---|---|---|---|
| Linear regression model variable | **Nusinersen** Estimate | **Risdiplam** Estimate | **Nusinersen** Estimate | **Risdiplam** Estimate |
| Intercept | 0.15296 | 0.17158 | 0.15297 | 0.17158 |
| SMN protein levels prior to treatment | 1.45840 | 1.50690 | 0.45840 | 0.50689 |
| *SMN2Δ7* mRNA levels prior to treatment | -0.10463 | -0.13511 | -0.10463 | -0.13511 |
| Age group (paediatric/adult) | -0.06415 | -0.05990 | -0.06415 | -0.05990 |
| *SMN2* copy number (3x *SMN2*) | 0.09871 | 0.11985 | 0.09871 | 0.11985 |
| *SMN2* copy number (4x *SMN2*) | 0.13355 | 0.17212 | 0.13355 | 0.17212 |

**Table S3. Linear regression model for SMN levels (total amount and increase) after treatment.** Estimates of the factors significant in the linear regression model for SMN protein increase after *in vitro* treatment.

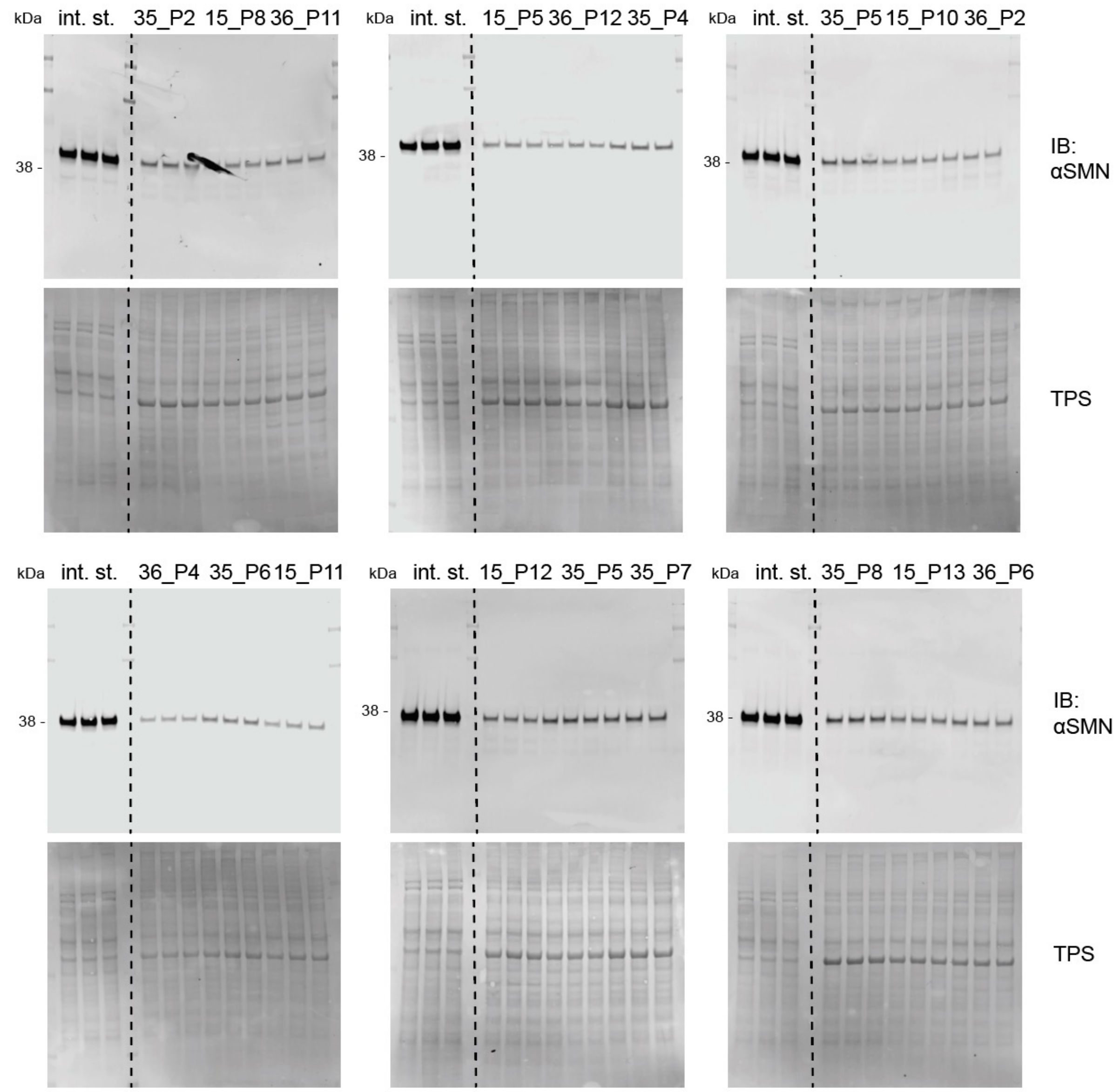


**Supplementary figure 8A**. Uncropped SMN western blots for Fig. S2 (SMN at different passage numbers). TPS = total protein staining; in. st = internal standard.

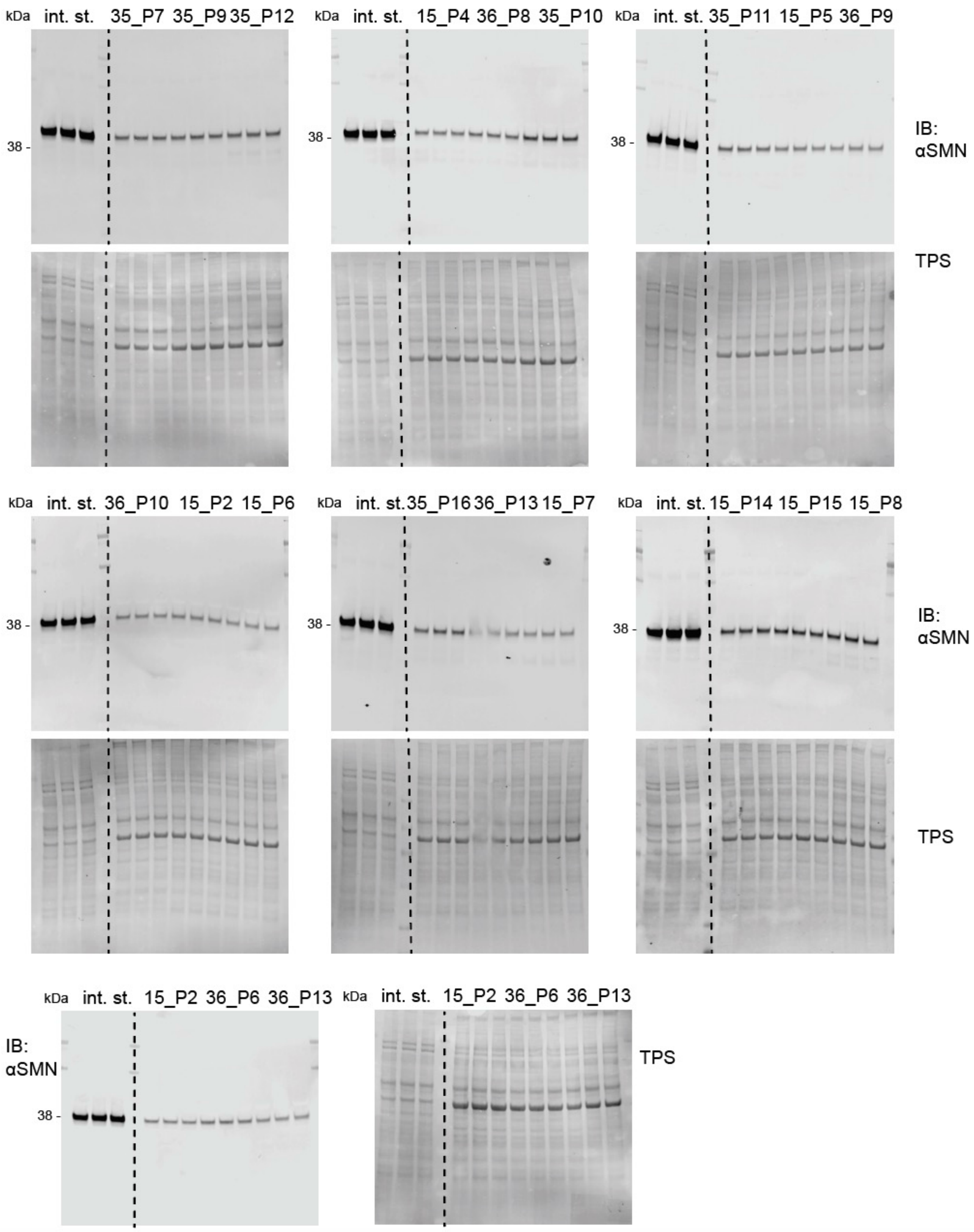


**Supplementary figure 8B**. Uncropped SMN western blots for Fig. S2 (SMN at different passage numbers). TPS = total protein staining; in. st = internal standard.

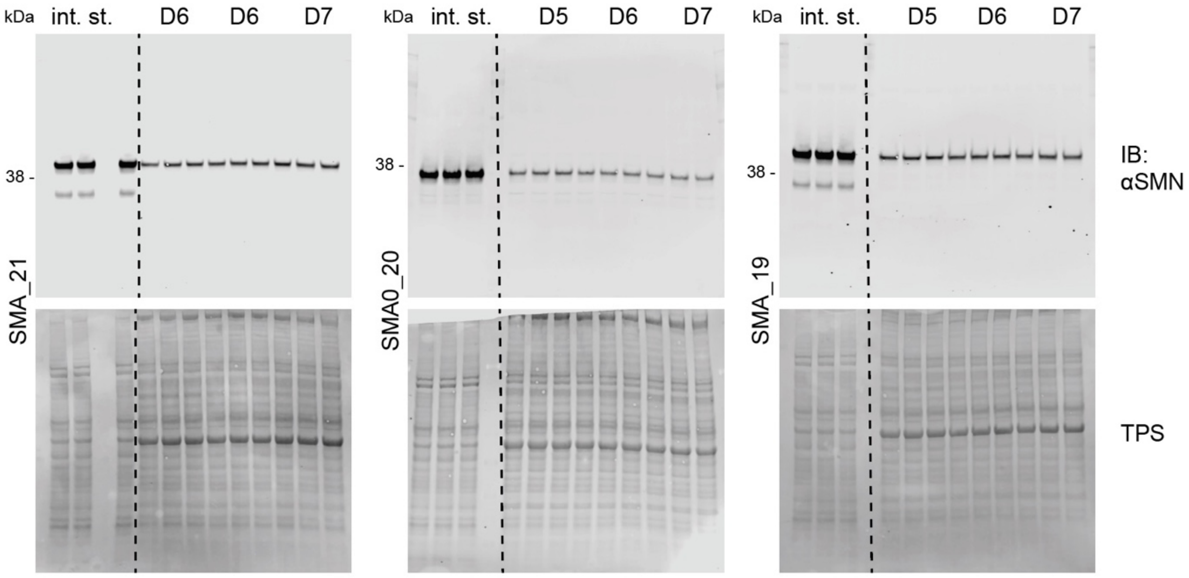


**Supplementary figure 8C**. Uncropped SMN western blots for Fig. S2 (SMN at different confluences and days in culture). TPS = total protein staining; in. st = internal standard.

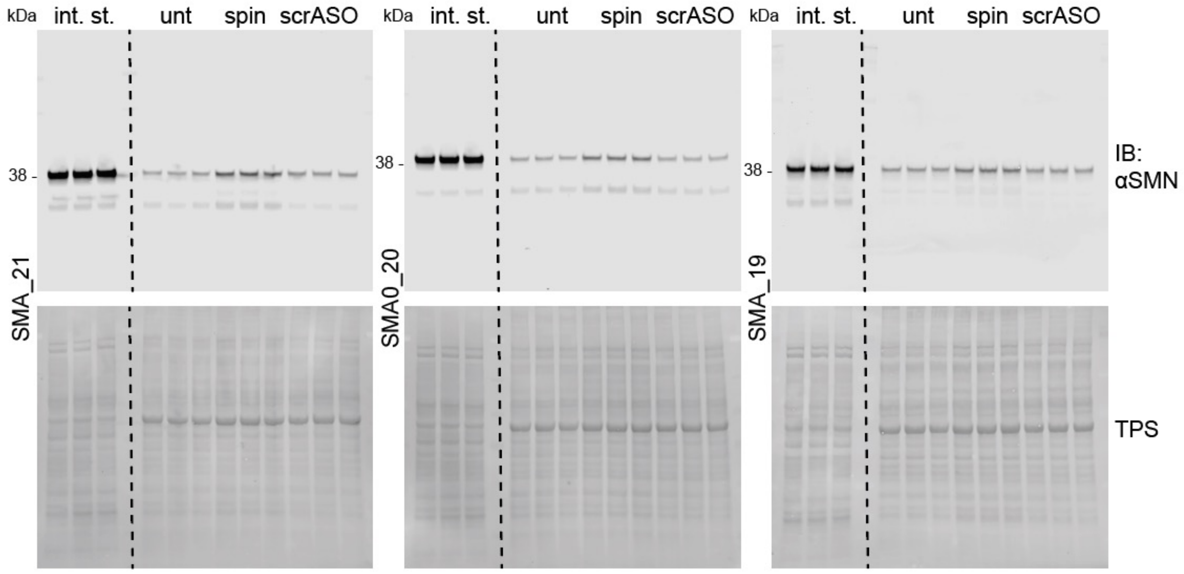


**Supplementary figure 8D**. Uncropped SMN western blots for Fig. S2. SMN protein levels in patient-derived fibroblasts untreated (U), treated with nusinersen (N) and treated with scrASO (scrASO). TPS = total protein staining; in. st = internal standard.

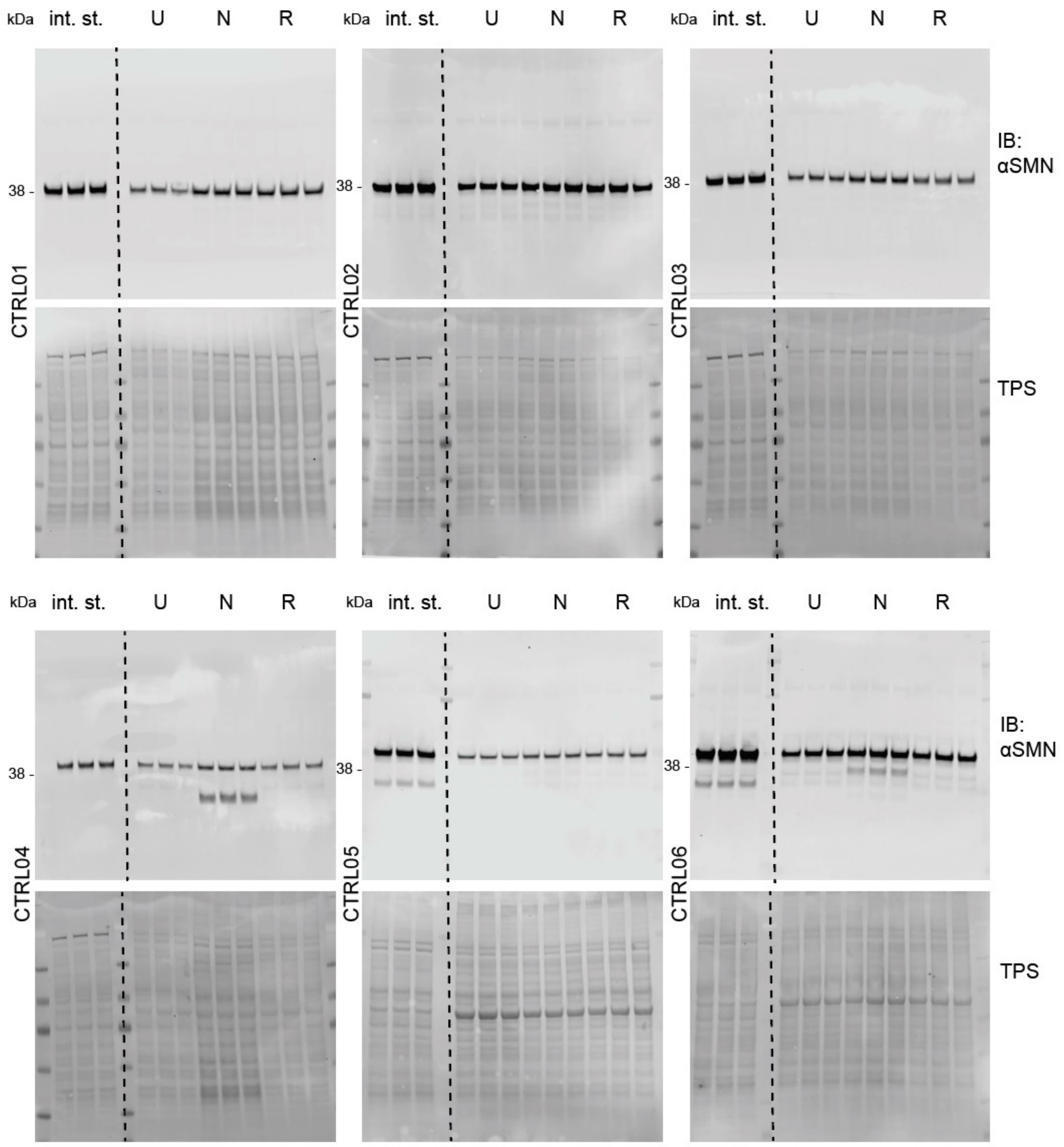


**Supplementary figure 8E**. Uncropped SMN western blots for Fig. 1, 2, 4, S5, S6, S7. SMN protein levels in patient-derived fibroblasts untreated (U), treated with nusinersen (N) and treated with risdiplam (R). TPS = total protein staining; in. st = internal standard.

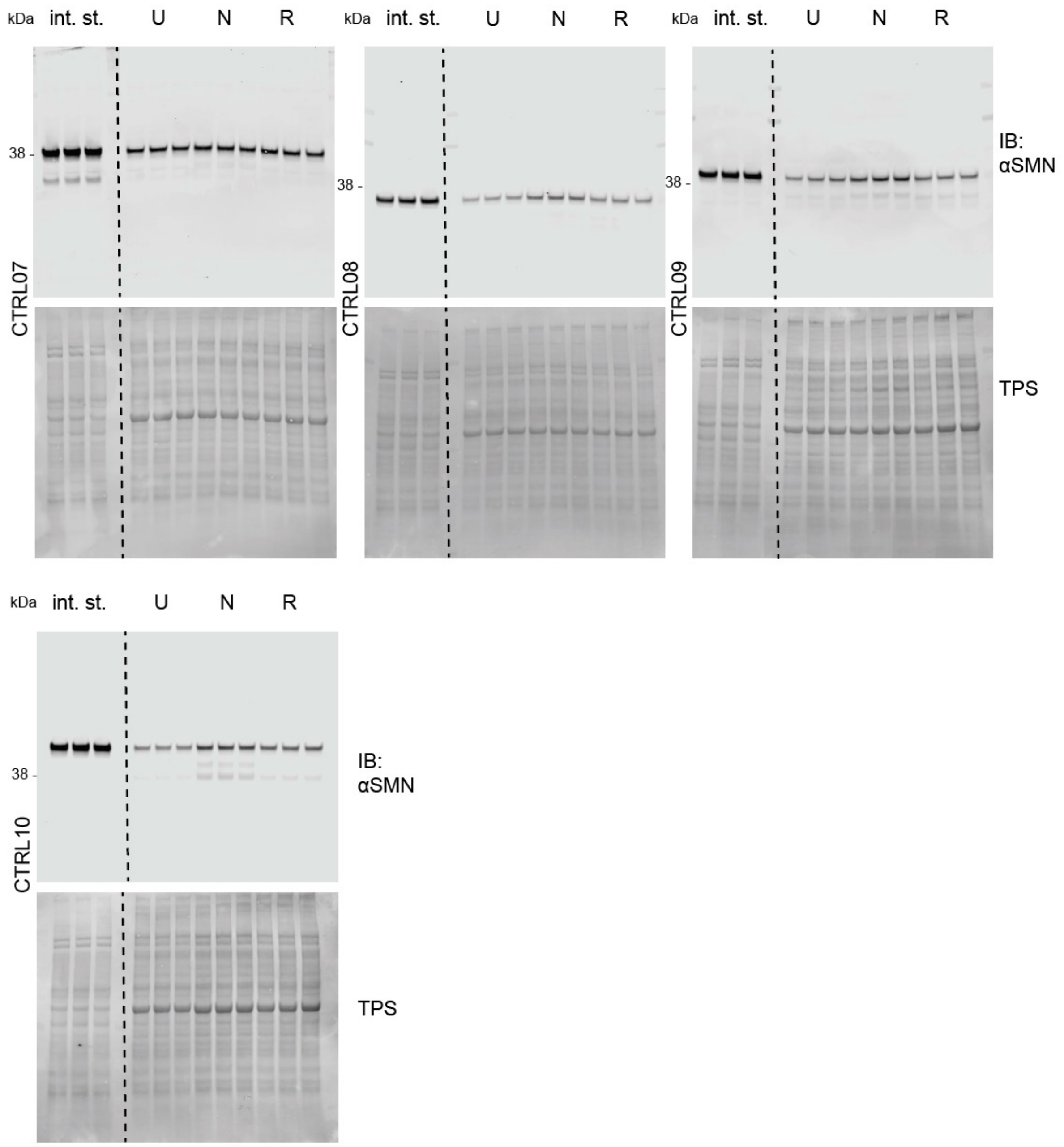


**Supplementary figure 8F**. Uncropped SMN western blots for Fig. 1, 2, 4, S5, S6, S7. SMN protein levels in patient-derived fibroblasts untreated (U), treated with nusinersen (N) and treated with risdiplam (R). TPS = total protein staining; in. st = internal standard.

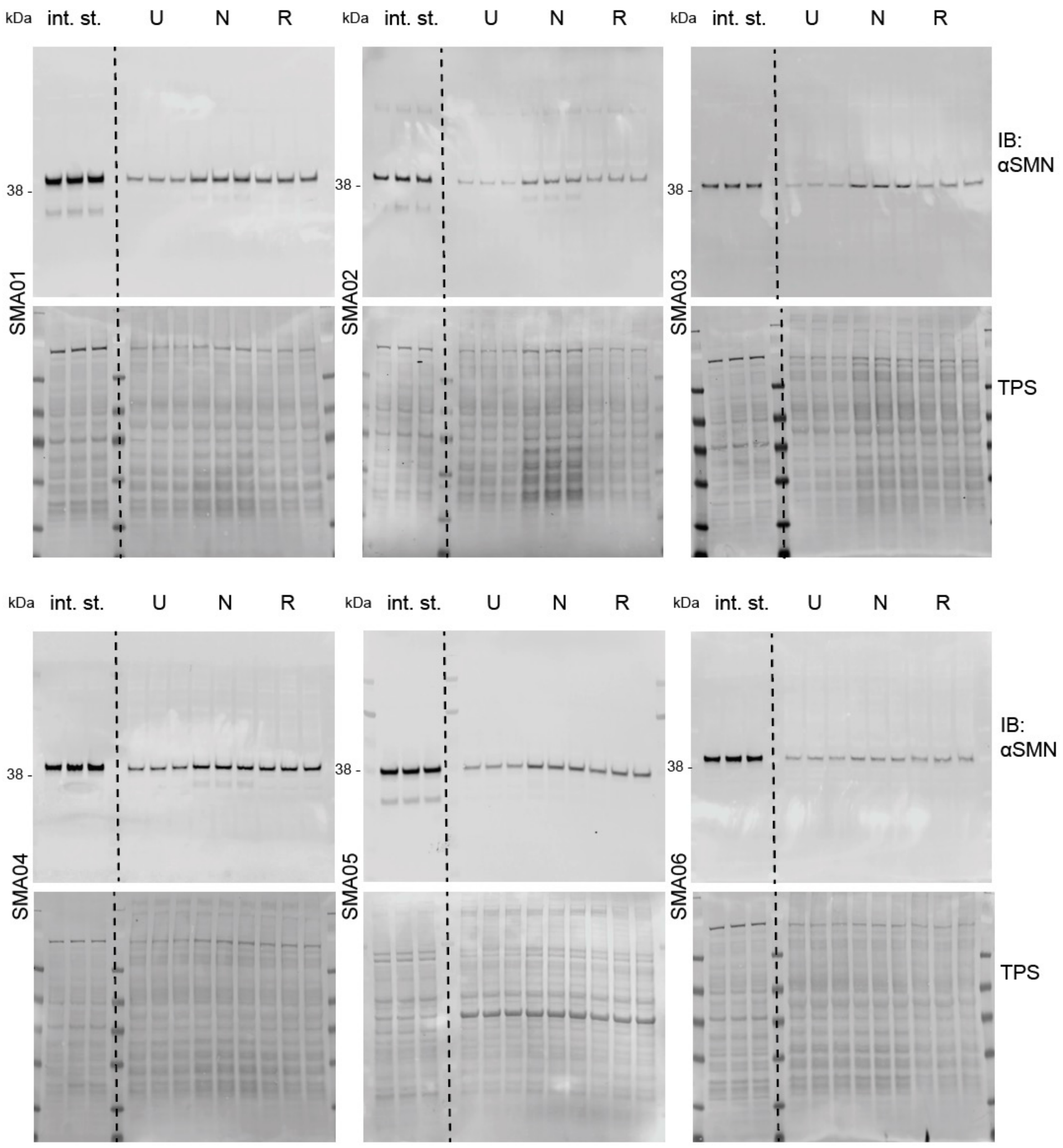


**Supplementary figure 8F**. Uncropped SMN western blots for Fig. 1, 2, 3, 4, S5, S6. SMN protein levels in patient-derived fibroblasts untreated (U), treated with nusinersen (N) and treated with risdiplam (R). TPS = total protein staining; in. st = internal standard.

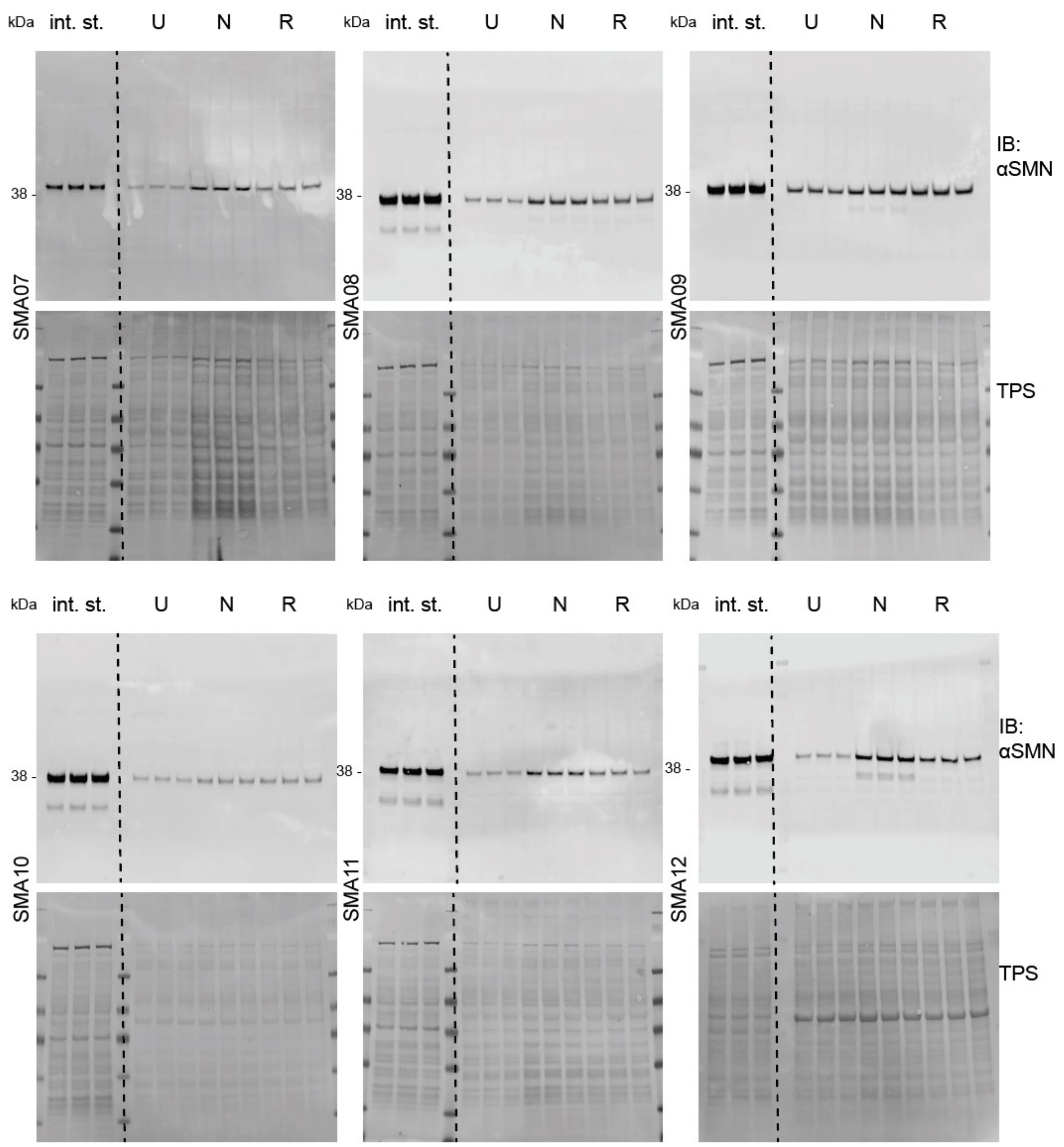


**Supplementary figure 8G**. Uncropped SMN western blots for Fig. 1, 2, 3, 4, S5, S6. SMN protein levels in patient-derived fibroblasts untreated (U), treated with nusinersen (N) and treated with risdiplam (R). TPS = total protein staining; in. st = internal standard.

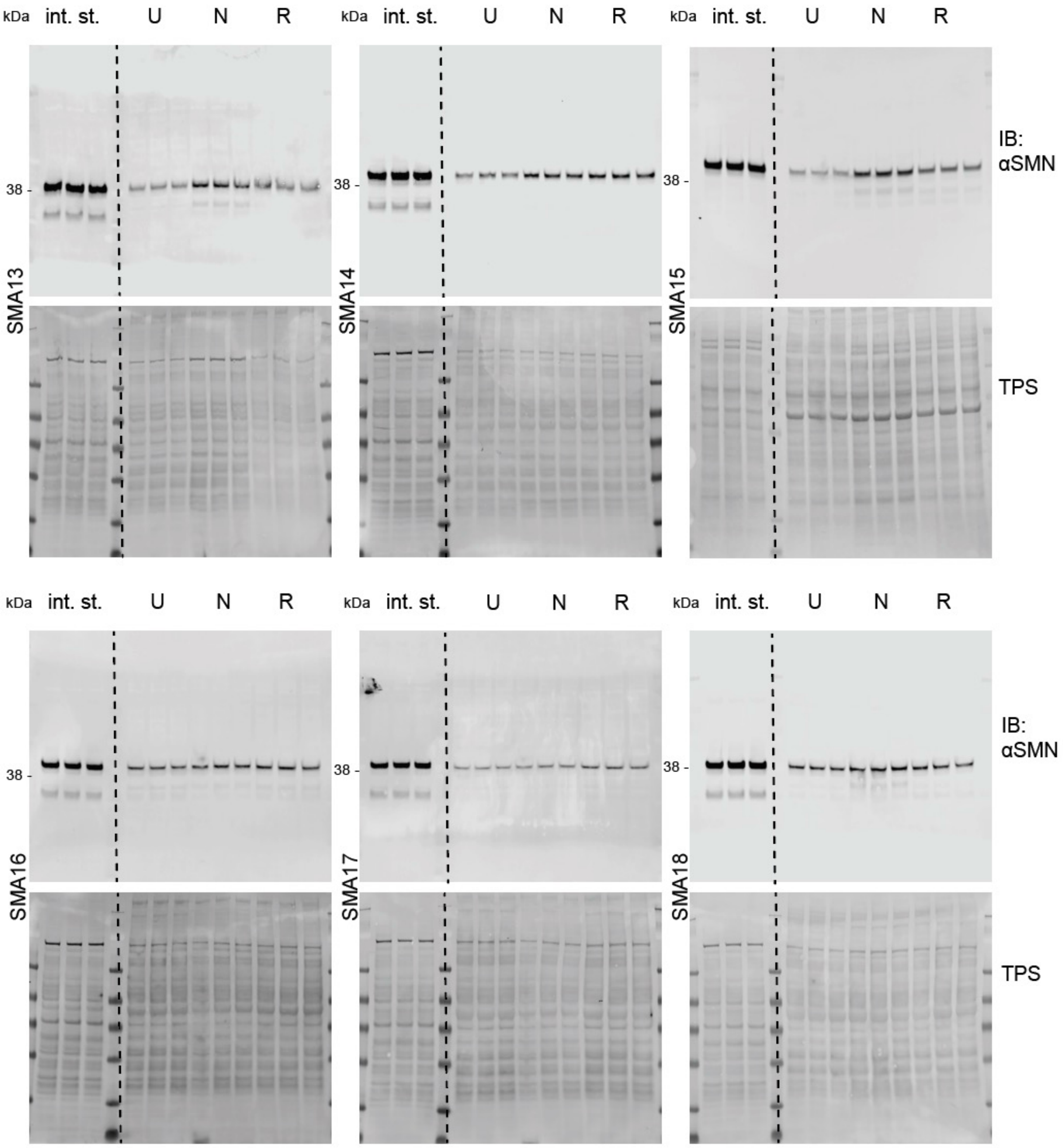


**Supplementary figure 8H**. Uncropped SMN western blots for Fig. 1, 2, 3, 4, S5, S6. SMN protein levels in patient-derived fibroblasts untreated (U), treated with nusinersen (N) and treated with risdiplam (R). TPS = total protein staining; in. st = internal standard.

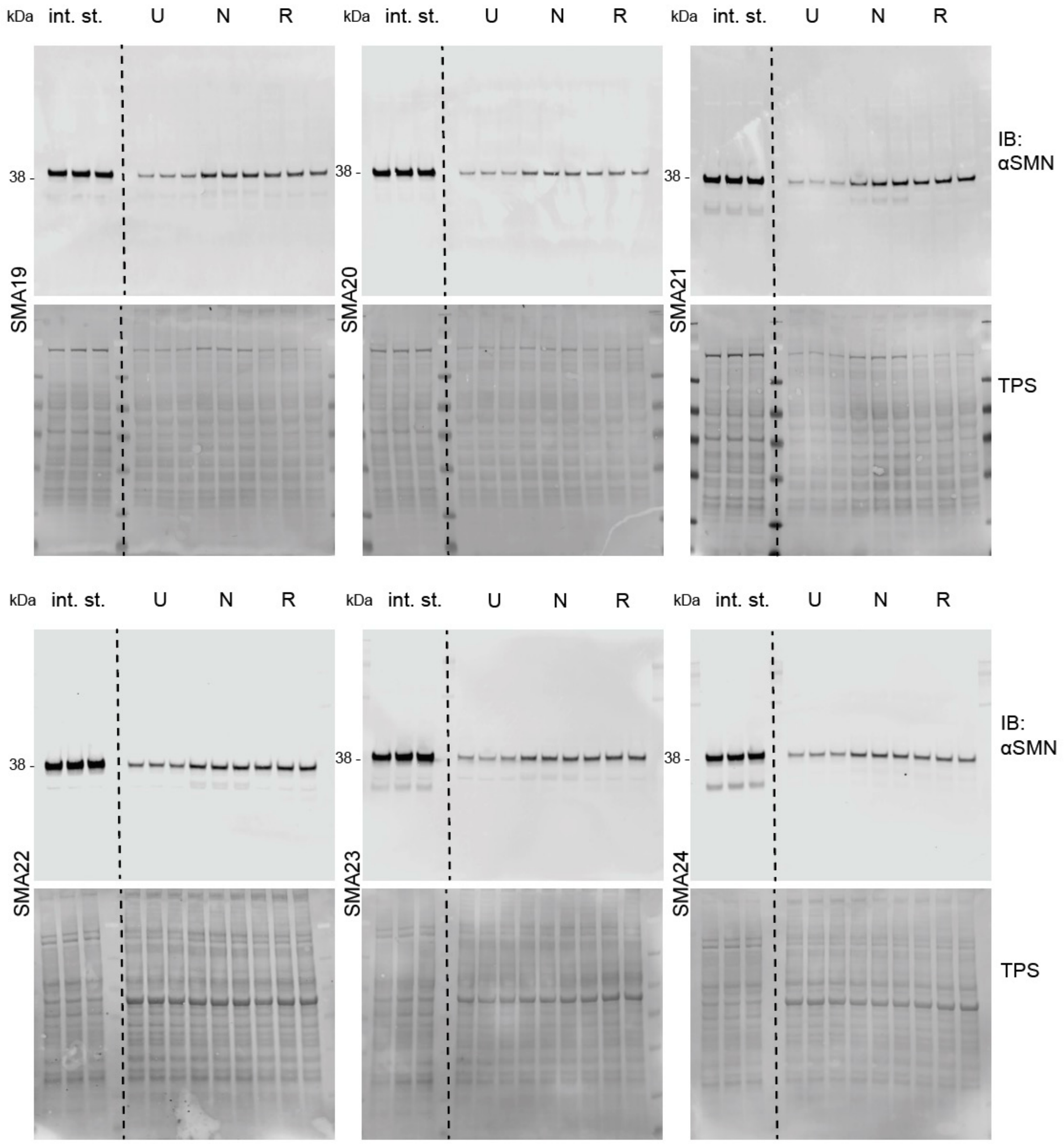


**Supplementary figure 8I.** Uncropped SMN western blots for Fig. 1, 2, 3, 4, S5, S6. SMN protein levels in patient-derived fibroblasts untreated (U), treated with nusinersen (N) and treated with risdiplam (R). TPS = total protein staining; in. st = internal standard.

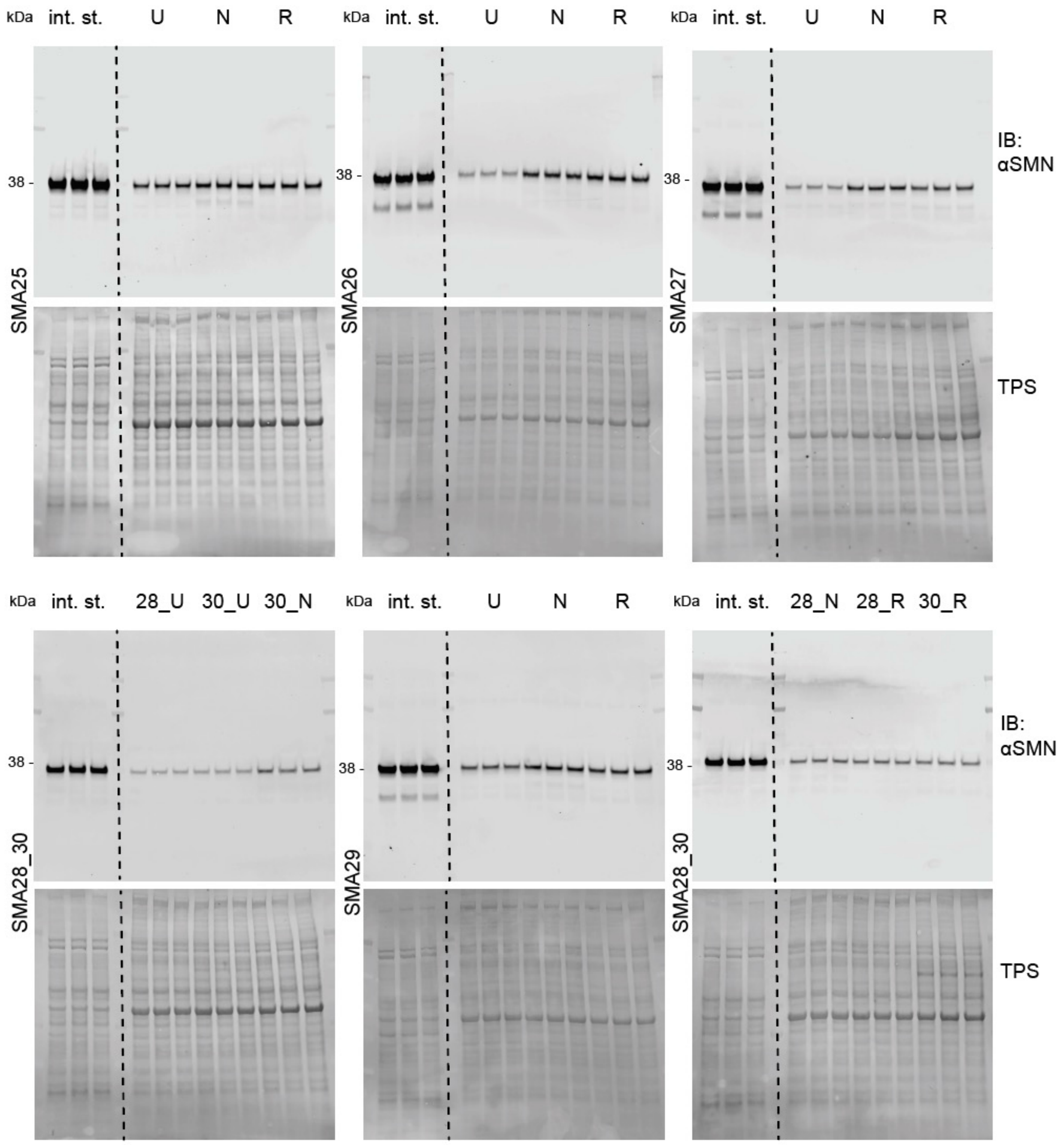


**Supplementary figure 8J**. Uncropped SMN western blots for Fig. 1, 2, 3, 4, S5, S6. SMN protein levels in patient-derived fibroblasts untreated (U), treated with nusinersen (N) and treated with risdiplam (R). TPS = total protein staining; in. st = internal standard.

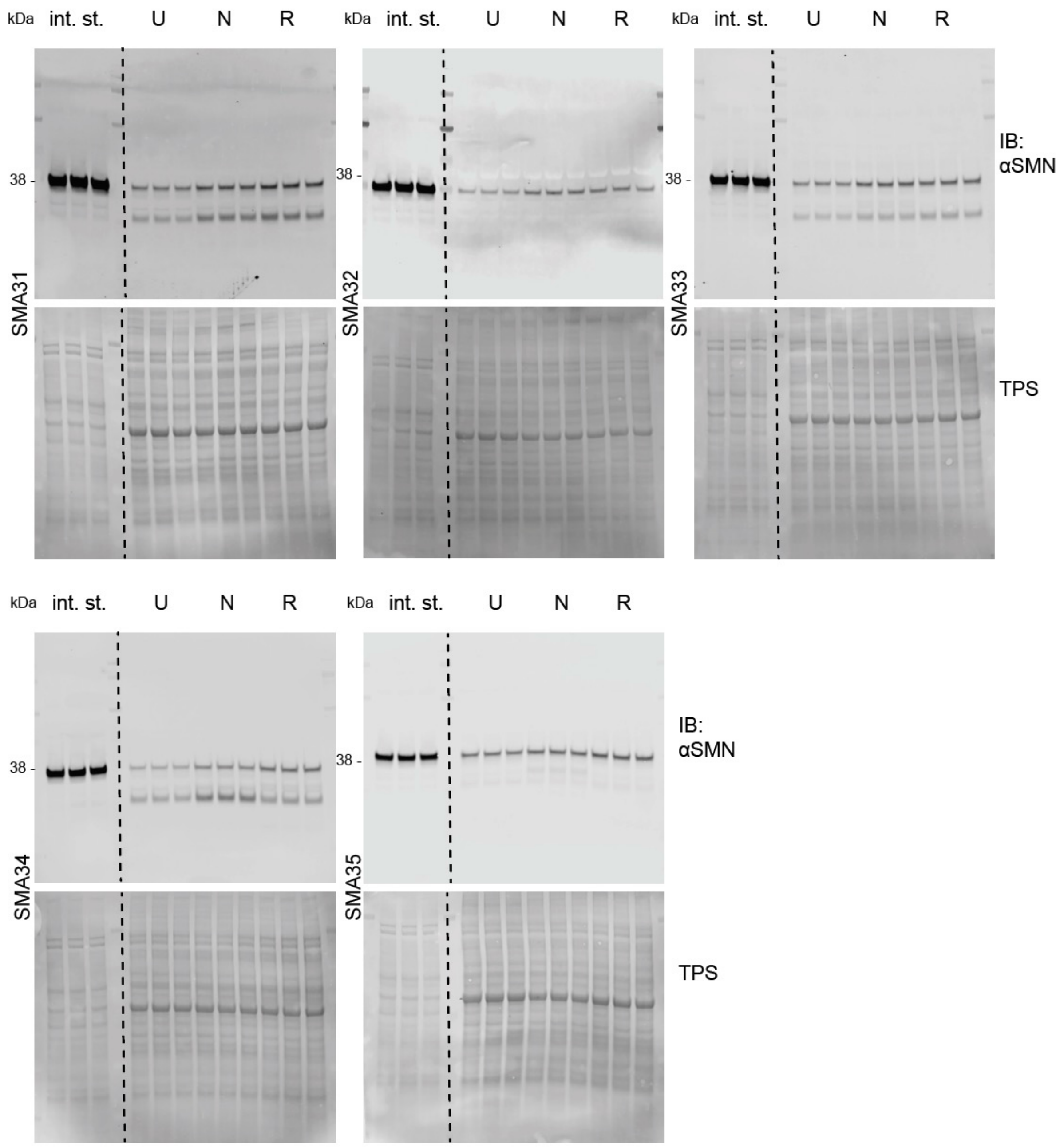


**Supplementary figure 8K**. Uncropped SMN western blots for Fig. 1, 2, 3, 4, S5, S6. SMN protein levels in patient-derived fibroblasts untreated (U), treated with nusinersen (N) and treated with risdiplam (R). TPS = total protein staining; in. st = internal standard.
